# PIFI Stabilizes Chloroplast NDH–PSI Supercomplex to Maintain Plastoquinone Redox Balance and PSII Efficiency

**DOI:** 10.64898/2026.03.22.713156

**Authors:** Kaori Kohzuma, Minami Murai, Ko Imaizumi, Kenta Miura, Ayaka Kimura, Keisuke Yoshida, Yufen Che, Noriko Ishikawa, Toru Hisabori, Kentaro Ifuku

## Abstract

Photosynthetic electron transport is mediated by several protein supercomplexes that are spatially arranged in the thylakoid membranes of chloroplasts. The chloroplast NADH dehydrogenase-like (NDH) complex is part of the photosynthetic alternative electron transport (AET) chain, which reduces the plastoquinone (PQ) pool using reduced ferredoxin as a substrate. This NDH complex is associated with photosystem I (PSI) and mediates a portion of AET in stroma lamellae, whereas photosystem II (PSII) is concentrated in grana stacks. This study presents the findings regarding <u>p</u>ost-illumination chlorophyll fluorescence increase (PIFI), a protein crucial for regulating AET via the NDH pathway. A marked increase in NDH activity and a reduction in the PQ pool in the dark were observed in *PIFI*-deficient mutant strains (*g-pifi*) generated by genome editing. Blue native PAGE analysis indicated that PIFI was associated with the NDH-PSI supercomplex in the wild type, and the NDH complex was dissociated from PSI in the *g-pifi* mutants. Additionally, the *g-pifi* mutants exhibited a decrease in the maximum quantum yield of PSII (Fv/Fm). Notably, Fv/Fm was restored in a double mutant harboring both *g-pifi* and NDH-deficient *pnsl1* mutations, demonstrating that deregulated NDH activity in *g-pifi* causes downregulation of PSII efficiency. However, the lower Fv/Fm was not observed in a mutant lacking thioredoxin m4 *(trxm4*), which showed deregulated NDH activity but maintained the NDH-PSI supercomplex. These data suggest that PIFI stabilizes the NDH-PSI supercomplex and maintains the spatial localization of PQ reduction via AET in thylakoid membranes, which is essential for the proper functioning of PSII.

## Introduction

In photosynthesis, light energy absorbed by the photosynthetic apparatus in the thylakoid membranes of chloroplasts is converted into chemical energy in the form of NADPH and ATP through electron transport reactions. These energy carriers drive metabolic pathways that fix CO_2_. The linear electron transport (LET) pathway transfers reducing capacity from water to NADPH via photosystem II (PSII), the plastoquinone (PQ) pool, cytochrome *b*_6_*f* complex (Cyt *b*_6_*f*), plastocyanin, photosystem I (PSI), ferredoxin (Fd), and Fd-NADP^+^-reductase (Johnson 2025). During this electron transport process, protons flow from the stroma into the thylakoid lumen, thereby establishing a proton gradient across the thylakoid membranes. This proton gradient drives ATP synthesis by the chloroplast ATP synthase (Witt 1979; Konno et al. 2006; Kohzuma et al. 2025). In addition to LET, there are alternative electron transport (AET) pathways that modulate the energy production required for photosynthesis (Alric and Johnson 2017).

AET refers to a set of electron transport pathways connected to LET, including cyclic electron transport around PSI (CET-PSI), the water-water cycle, photorespiration, and chlororespiration (Bennoun 1982; Miyake and Asada 1992; Munekage et al. 2002; Tikkanen and Aro 2026). AET helps dissipate this excess reducing power and maintain a balanced redox state of the electron transport chain: Excessive light exposure can over-reduce the electron transport chain, leading to the formation of reactive oxygen species (ROS) and photoinhibition (Asada 1999; Krieger-Liszkay and Shimakawa 2022, Sonoike 2025; Takeuchi et al. 2025). The accumulation of reducing power poses a risk not only under high-light conditions but also during dark-to-light transitions; if light exposure occurs suddenly when the PQ pool is over-reduced, it can cause photoinhibition and damage to the photosynthetic machinery (Kono and Terashima 2014; Tikhonov et al. 2015). It has also been reported that the reduction of the PQ pool under dark-chilling conditions, which triggers state transitions, can interfere with the proper localization of light-harvesting complex (LHC) II (Krysiak et al. 2024). To regulate the redox state of the PQ pool, plastid terminal oxidase (PTOX) directly oxidizes PQ and participates in chlorororespiration (Joët et al.2002; Rog et al. 2022; Messant et al. 2024). In addition, CET-PSI, which transfers electrons cyclically from Fd on the electron acceptor side of PSI back to the PQ pool, requires regulation to prevent negative effects on photosynthetic reactions (Johnson 2011; Suorsa 2015; Yamori and Shikanai 2016; Shikanai and Yamamoto 2017; Satoh et al. 2025).

The major function of CET-PSI has been proposed to be the regulation of the ATP/NADPH production ratio in photosynthetic electron transport, as it can generate a proton gradient without producing NADPH (Munekage et al. 2004; Zhang et al. 2023). There are two known pathways for CET-PSI: the PGR pathway, which depends on proton gradient regulation (PGR) proteins, and the NDH pathway, which involves the chloroplast NADH dehydrogenase-like (NDH) complex (Shikanai et al. 1998; Burrows et al. 1998; Munakage et al. 2002; Dal Corso et al. 2008). PGR5 is a key factor required for efficient cyclic electron flow in the PGR-dependent pathway, whereas the precise molecular components responsible for PQ reduction in this route remain under discussion (Rühle et al. 2021; Degen et al. 2023). The contribution of the NDH pathway to PSI-CET is considered low, especially in model plants, because of the low abundance of the NDH complex, which constitutes only a small percentage of PSI or PGRL1 (Burrows et al. 1998; Hertle et al. 2013). However, an increasing number of studies have suggested the physiological significance of the NDH pathway in C_4_ plants and plants under stressed conditions (Takabayashi et al. 2005; Ishikawa et al. 2016; Yamori et al. 2016; Ogawa et al. 2023; Ermakova et al. 2024; Takeuchi et al. 2025).

The chloroplast NDH complex is a large membrane protein complex comprising five subcomplexes, consisting of subunits derived from over 30 genes encoded by the chloroplast and nuclear genomes (Ifuku et al. 2011; Shikanai 2016; Kato et al. 2021; Shikanai 2025). The NDH complex interacts with the PSI complex to form a supercomplex, in which multiple PSI complexes could bind to a single NDH complex (Yadav et al. 2017). The most prevalent of these configurations is the NDH-PSI supercomplex, in which two PSI complexes are bound on either side of the NDH complex (Peng et al. 2009; Kouřil et al. 2014). These two PSI complexes, referred to as right-PSI and left-PSI, structurally stabilize the NDH complex by sandwiching it (Shen et al. 2022; Su et al. 2022; Introini et al. 2025). The PSI complex comprises a reaction center complex containing PsaA and PsaB, as well as four light-harvesting complex I (LHCI) proteins. LHCI proteins form heterodimers in which Lhca1-Lhca4 and Lhca2-Lhca3 are paired together (Ben-Shem et al. 2003; Pan et al. 2018). When PSI forms a supercomplex with NDH, Lhca4 is replaced by Lhca5 in the left PSI, and Lhca2 is replaced by Lhca6 in the right PSI, resulting in the formation of Lhca1-Lhca5 and Lhca6-Lhca3 heterodimers, respectively (Peng et al. 2009; Kouril et al. 2014; Otani et al. 2018; Shen et al. 2022; Su et al. 2022; Introini et al. 2025). The association of Lhca5 and Lhca6 with PSI stabilizes the NDH complex, which may facilitate efficient electron transfer from PSI via Fd, thereby promoting CET-PSI (Kato et al. 2018; Peng and Shikanai 2011).

The <u>p</u>ost-illumination chlorophyll fluorescence increase (PIFI) protein in *Arabidopsis thaliana* has been implicated in the regulation of PSI-CET via the NDH pathway (Wang and Portis 2007). Initially, PIFI attracted attention for its sequence similarity to a portion of the RuBisCO activase and for its classification as a stromal protein. In the T-DNA insertion mutant strain *pifi* (SALK_085656), NDH activity is impaired, as indicated by a decrease in post-illumination chlorophyll (Chl) fluorescence (Wang and Portis 2007). The maximum quantum yield of PSII (Fv/Fm), a measure of PSII efficiency, also decreased in the *pifi* mutant, especially under high light intensity, whereas other NDH-deficient mutants, such as *crr2* and *crr6*, did not exhibit declines in Fv/Fm (Hashimoto et al. 2003; Munshi et al. 2006). An additional feature of PIFI is its remarkably rapid turnover, comparable to that of the PSII reaction center protein D1 (Li et al. 2017), and it has been reported to undergo phosphorylation (Wang et al. 2013). Nevertheless, the precise function of PIFI within the NDH complex and the mechanism by which its absence affects PSII remain unclear.

This study revealed an unexpected deviation from previous findings, showing that the loss of PIFI leads to an increase in NDH activity. Genome-editing technology was used to create two *PIFI*-deficient mutant strains (*g-pifi_1* and *g-pifi_2*) that did not show decreased NDH activity, but instead exhibited increased activity and a reduced PQ pool. Consistent with previous reports, a decrease in Fv/Fm was observed in the new *g-pifi* mutants. The NDH complex appeared to dissociate from PSI in the *g-pifi* mutants. Notably, the Fv/Fm value was restored in the double mutant produced by crossing *g-pifi_1* with *pnsl1*, a *PNSL1*-deficient mutant that is a lumenal subcomponent of the NDH complex (Ishihara et al. 2007; Ifuku et al. 2011). These findings suggest that PIFI plays a vital role in regulating NDH activity to maintain the proper redox balance of the PQ pool in the dark and optimize PSII function.

## Results

### Generation of *PIFI*-deficient mutants using genome editing and the phenotypes of high NDH activity

In this study, we generated two new *PIFI*-deficient mutants (*g-pifi_1* and *g-pifi_2*) using CRISPR-Cas9 genome editing (Figure 1A, Supplemental Figure S1) (Tsutsui and Higashiyama 2017), in addition to the previously reported *PIFI*-deficient *pifi* strain of SALK_085656 carrying a T-DNA insertion in an intron of the *PIFI* gene (Wang and Portis 2007). Because the newly generated genome-edited strains contain mutations introduced into the exons, the signals detected in the immunoblot for the PIFI protein in those mutants are likely due to background (Figure 1B). No obvious changes were observed in the accumulation of subunit proteins of other representative components of PSII and PSI. To examine the relationship between PIFI and the NDH pathway of CET-PSI, we analyzed the *pnsl1* and *g-pifi_1* × *pnsl1* double mutant (Figure 1A) (Ishihara et al. 2007). PIFI accumulation was lowered to approximately 50% of the wild type in *pnsl1* and was undetectable in the *g-pifi_1* × *pnsl1* double mutant.

**Figure 1.**
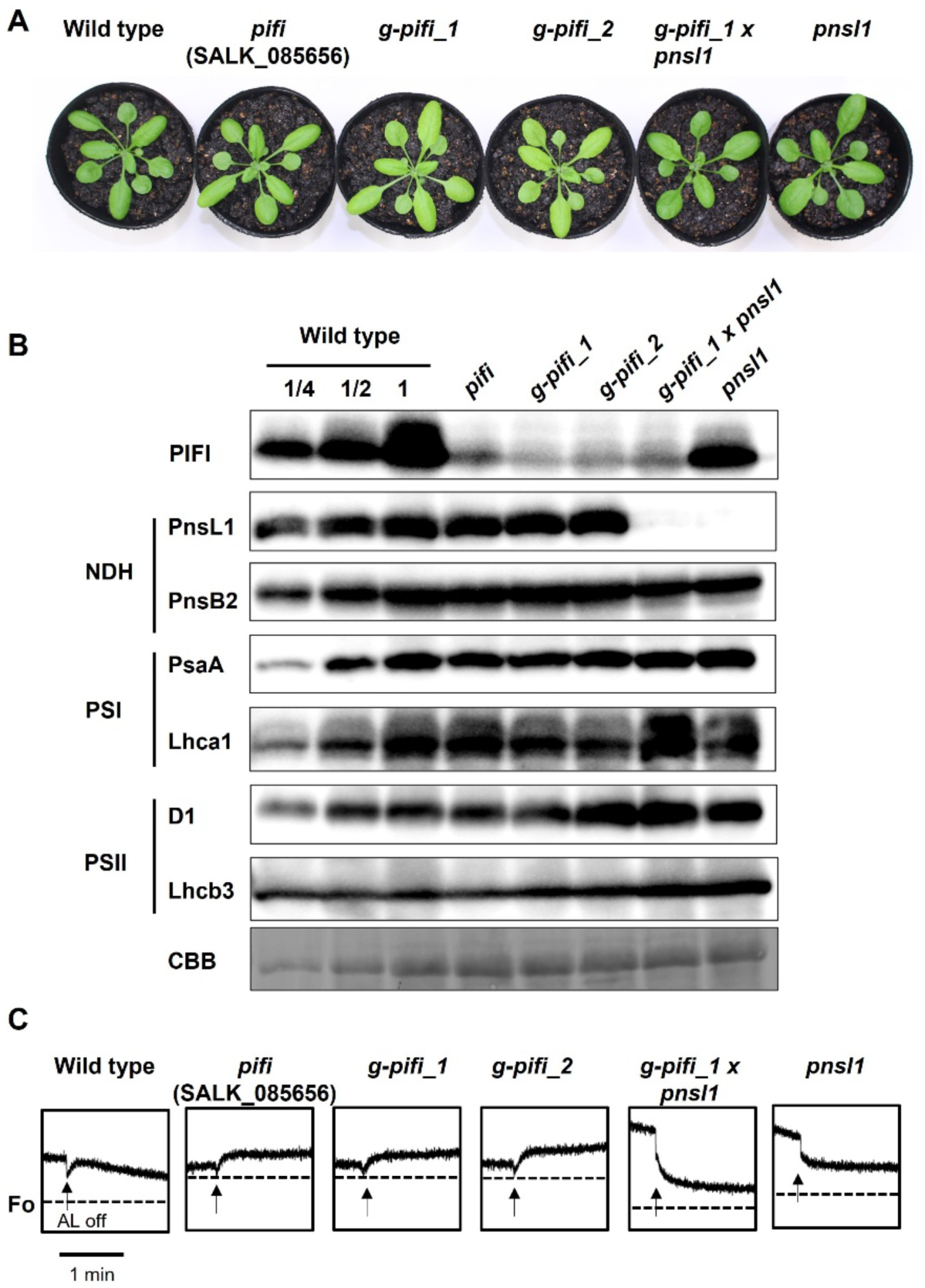
Phenotypes of wild type (Col-0), T-DNA inserted *pifi*, two strains of genome-edited *pifi* (*g-pifi_1*, *g-pifi_2*), and *g-pifi_1* crossed with *pnsl1*, an NDH-deficient mutant. A. Photographs of the plants grown for 3 weeks under long-day conditions. B. Immunoblot analysis using whole-leaf proteins. Whole-leaf proteins were isolated from wild-type (Col-0) and mutants. Proteins were separated using SDS-PAGE and immunodetected using specific antibodies against the PIFI, NDH complex (PnsL1 and PnsB2), PSI (PsaA and Lhca1), and PSII (D1 and Lhcb3). For the wild type, a dilution series of proteins corresponding to 20.0 (100%), 10.0, and 5.0 µg of whole proteins was loaded. Other mutants contained whole proteins corresponding to 2.0 µg in each lane. The accumulation level in the CBB-stained gel is shown as a control. C. Post-illumination Chl *a* fluorescence used to evaluate NDH activity. The trace represents 1.5 minutes after turning off the AL. The dotted line indicates Fo. Bars = 1 min.

The NDH activities of the above mutants were then monitored by observing the transient increase in Chl fluorescence after switching off actinic light (AL) (Shikanai et al. 1998) (Figure 1C, Supplemental Figure S3). This transient increase reflects the re-reduction of the PQ pool in the dark via the NDH pathway of CET-PSI, accompanied by reduced stromal Fd. Previous studies have shown that the *pnsl1* mutant, does not exhibit increased Chl fluorescence upon AL inactivation (Ishihara et al. 2007). In contrast, the *pifi* mutant (SALK_085656) and two *g-pifi* mutants showed a continuous increase in Chl fluorescence after AL was turned off, unlike the wild-type and *pnsl1* mutants. This observation contrasts with previous reports (Wang and Porits 2007). Furthermore, in the *g-pifi_1* × *pnsl1* double mutants, this increase in Chl fluorescence was not observed, indicating that it was due to NDH activity.

To clarify the cause of the discrepancies between the present and previous results, we investigate several possible factors using the plants at various growth stages. Among these, we found that the NDH activity in *g-pifi_1* was significantly affected by the leaf developmental stage (Supplemental Figure S4). When we analyzed a 3-week-old plant, we observed an increase in NDH activity in relatively young leaves (around the 10th leaf in *g-pifi_1*) immediately after full expansion. However, a loss of NDH activity was observed in the relatively senescent leaves around the 5th leaf in *g-pifi_1* plant, whereas PSII activity remained essentially unchanged. In contrast, in the wild-type, no significant difference in NDH activity was observed between the upper and lower leaves. This suggests that NDH activity becomes unstable during leaf development in *PIFI*-deficient mutants.

It should be noted that all three *PIFI*-deficient mutant strains (*pifi*, *g-pifi_1*, and *g-pifi_2*) showed slightly paler green leaves than the wild-type (Figure 1A). Consistently, their Chl levels were significantly lower than those of the wild type with decreased light-harvesting capacity, and the *g-pifi_1* × *pnsl1* double mutant showed recovery in Chl content (Supplemental Figure S2). This suggests that the pale-green phenotype of the *PIFI*-deficient mutants is linked with NDH activity.

### PQ pool reduction in *PIFI*-deficient mutants

An increase in NDH activity is presumed to induce a reduction in the PQ pool. To verify that the continuous increase in Chl fluorescence in the *pifi* and *g-pifi* mutants was due to a reduction in the PQ pool, far-red (FR) light was applied during the measurement of NDH activity (Figure 2A). FR irradiation specifically excites PSI, thereby oxidizing the PQ pool. In the wild type, Chl fluorescence transiently increased after AL was switched off and then decreased, but further decreased when FR was irradiated. In *pnsl1*, the Chl fluorescence level did not decrease when FR was applied because electrons did not flow into the PQ pool via NDH. In the *pifi* and *g-pifi* mutants, the fluorescence levels decreased significantly during FR irradiation. These results suggest that the increased Chl fluorescence observed in the *pifi* and *g-pifi* mutants was due to a reduction in the PQ pool. Furthermore, in the *g-pifi_1* × *pnsl1* double mutant, the increase in Chl fluorescence was abolished, showing a phenotype similar to *pnsl1*. This strongly suggests that PIFI is involved in the redox regulation of the PQ pool via the NDH pathway.

**Figure 2.**
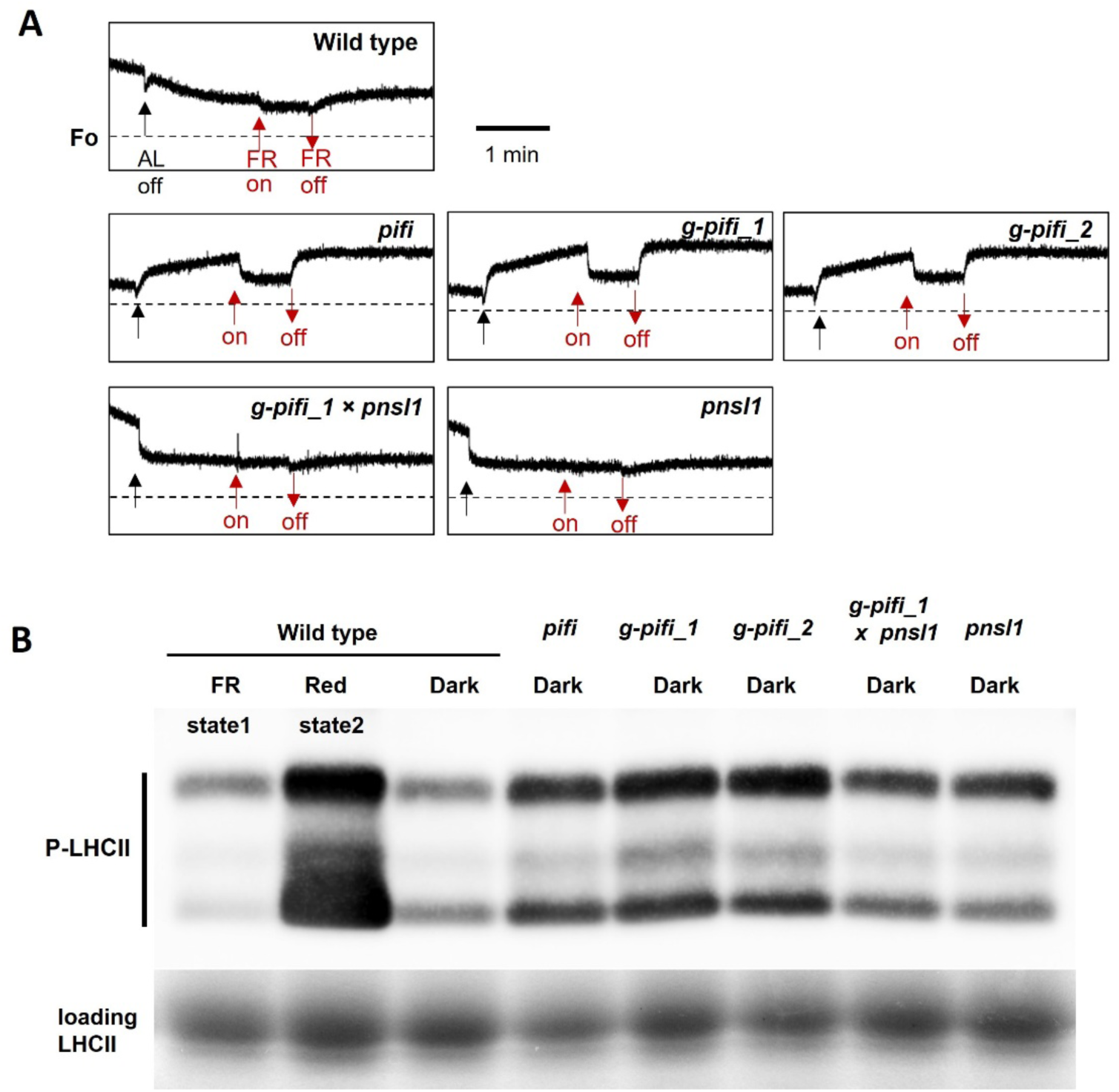
Changes in NDH activity and the redox state of the PQ pool. A. Evaluation of PQ pool reduction by Far-Red exposure in darkness. The black arrow indicates the point at which the actinic light (AL) is turned off. The red arrows indicate the points at which the far-red (FR) light is turned on and off. The dotted line indicates Fo. Bars = 1 min. B. Phosphorylation state of LHCII under dark. Thylakoid membranes were isolated from wild-type (Col-0) and mutant plants grown in the dark. Proteins were separated using SDS-PAGE and immunoblotted with an anti-phospho-threonine antibody to detect phosphorylated LHCII (P-LHCII). In the wild type, LHCII was dephosphorylated under far-red light for 10 min (state 1) and phosphorylated under red light for 10 min (state 2), serving as negative and positive controls, respectively. The lower panel shows total LHCII as a loading control.

A reduction in the PQ pool induces LHCII phosphorylation (Nellaepalli et al. 2012). The phosphorylation state of LHCII after 2 h of dark adaptation was analyzed using an anti-phosphorene antibody (Figure 2B). As controls, wild-type leaves were exposed to FR light (730 nm) to oxidize the PQ pool or to red light (red, 660 nm) to reduce it, thereby inducing dephosphorylation (state 1) and phosphorylation (state 2) conditions, respectively. In dark-treated wild-type leaves, LHCII was dephosphorylated and showed a weak signal, whereas all thr *pifi* and *g-pifi* mutants exhibited stronger phosphorylation signals than the wild type. In contrast, the *pnsl1* and *g-pifi_1* × *pnsl1* mutants without NDH activity did not exhibit increased LHCII phosphorylation. These results indicated that the *pifi* and *g-pifi* mutants with increased NDH activity exhibited a reduced PQ pool in the dark.

### PIFI stabilizes the NDH-PSI complex

To investigate the mechanism by which the loss of PIFI enhances CET in the NDH pathway and promotes the reduction of the PQ pool, thylakoid membrane protein complexes from wild-type, *g-pifi_1*, *g-pifi_2*, and *g-pifi_1* × *pnsl1* were separated using BN-PAGE (Figure 3A). The bands that were significantly altered in the *g-pifi_1* mutant compared to those in the wild-type were excised and analyzed using LC-MS/MS (Figure 3B, Supplemental Table 1). Band I contained PSI-LHCI and NDH complexes, representing the NDH-PSI_2_ supercomplex. In *g-pifi* mutants, the accumulation of Band I decreased, whereas two new bands appeared in the Band II region (black arrows) (Figure 3A). Bands I and II were not observed in *g-pif_1* × *pnsl1*, indicating that they included the NDH complex. Band II contained PSI and NDH subunits, including NdhF, PnsB1, PnsL1, Lhca5, and Lhca6, whereas NdhH and NdhU were not detected. In addition, Band III showed a significant increase in *g-pifi* mutants and contained PSI, which would dissociate from the NDH-PSI_2_ supercomplex. All of the Band I–III, in particular Band III, contain ATP synthase α and β subunits, but their levels did not differ among the samples (Figure 3B). These results suggest that in *g-pifi* mutants, the NDH-PSI_2_ supercomplex is unstable, leading to the dissociation of the NDH complex and PSI core, generating Bands II and III (Figure 3C). This observation was also supported by BN-PAGE followed by SDS-PAGE in the second dimension (2D BN/SDS-PAGE) and immunoblotting with specific antibodies (Supplemental Figure S5).

**Figure 3.**
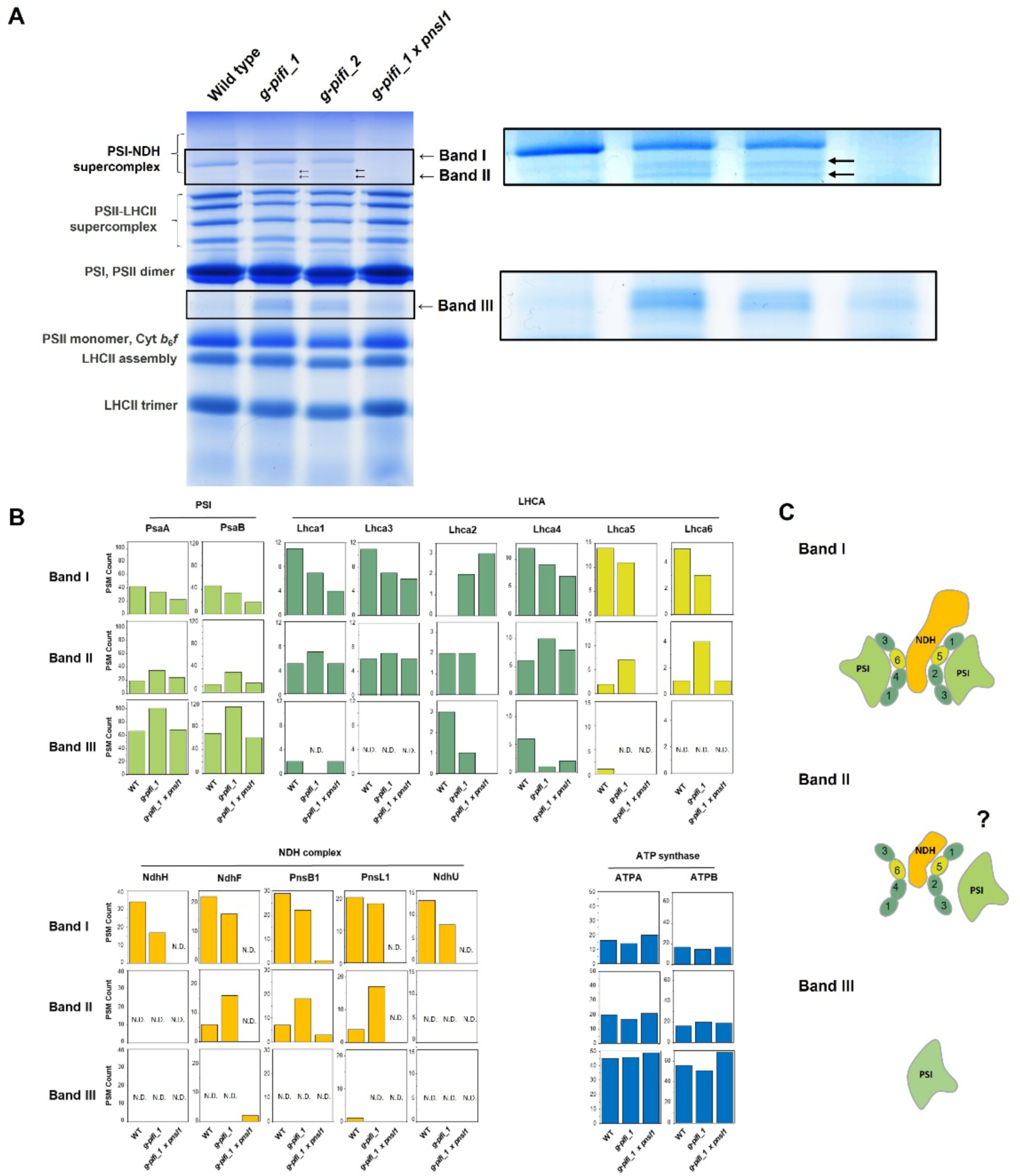
Comparative analysis of the formation of the PSI-NDH supercomplex in wild type and *PIFI*-deficient mutants. A. Principal photosynthesis protein complexes were separated using BN-PAGE. Thylakoids were isolated from the leaves of wild-type (Col-0) and mutant plants, and then solubilized with 1.0% (w/v) n-dodecyl-β-D-maltoside. Proteins corresponding to 10 µg of Chl were separated on a 3–12% gradient gel. The enlarged images are displayed on the right. B. The significantly changed bands were sliced as Band I, Band II, and Band III, and these contents were identified using LC-MS/MS. The number of peptides in representative components of the PSI core, NDH complex, and ATP synthase was compared using bar graphs. Full results are shown in Supplemental Table S1. C. Components of the PSI-NDH supercomplex suggested by mass spectroscopy and immunoblot analysis. N.D., not detected.

In the MS analysis, PIFI peptides were detected in Band I in the wild type (WT), which corresponds to the NDH-PSI_2_ supercomplex (Table S1). Immunoblotting with a PIFI-specific antibody detected bands in Band I and at a lower molecular mass region in the WT (Figure 4A). SDS-PAGE indicated that this signal was approximately 25 kDa, which corresponded to the mature PIFI protein. Furthermore, no signal was observed for *g-pifi_1*. These results suggest that PIFI interacts with the NDH-PSI supercomplex, although it may also exist independently. PIFI was previously reported to localize in the stroma; therefore, we re-examined its subchloroplast localization by fractionating isolated chloroplasts into stromal and thylakoid fractions and performing an immunoblot analysis (Figure 4B). These results clearly indicated that PIFI was predominantly localized in the thylakoid membrane fraction.

**Figure 4.**
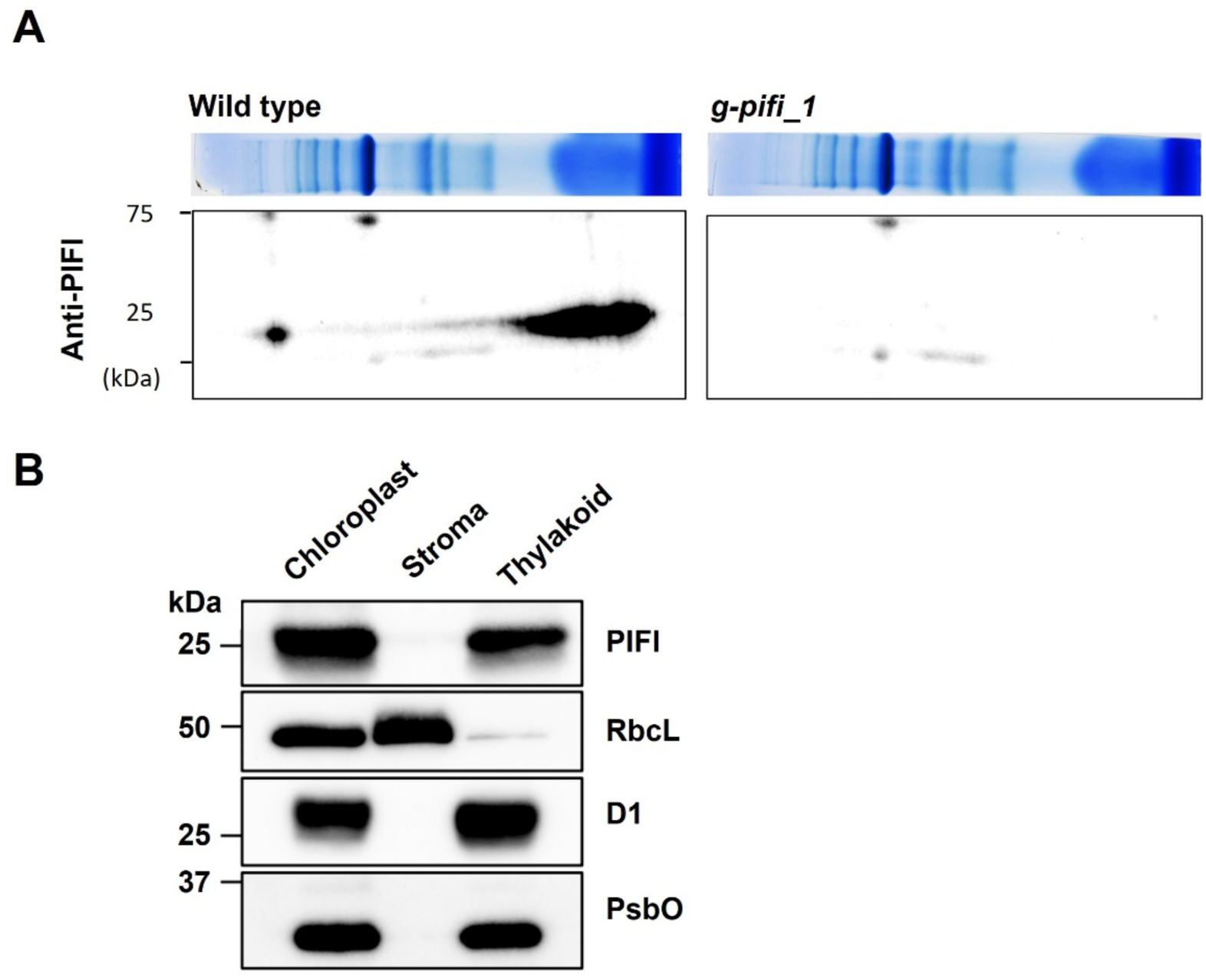
Localization of PIFI. A. Protein complexes separated using BN-PAGE were subsequently subjected to 2D SDS-PAGE in wild-type and *g-pifi_1*. PIFI was detected using specific antibodies raised against each protein. B. Immunoblot analysis to determine the subchloroplast localization of PIFI. Chloroplasts were fractionated into total, stroma, and thylakoid fractions. PIFI was predominantly detected in the thylakoid fraction. RbcL was used as a marker for the stroma, while D1 and PsbO served as markers for the thylakoids.

### Lower Fv/Fm in *PIFI*-deficient mutants is due to abnormal PQ reduction via NDH activity

Another distinguishing feature of *pifi* and *g-pifi* mutants was their lower Fv/Fm values than those of the WT (Figure 5A). This correlates with the pale-green leaf color in mutants with lower Chl accumulation (Figure 1A). The Fv/Fm value of *pnsl1* was similar to that of the wild-type, suggesting that the absence of the NDH complex does not impair PSII efficiency. Interestingly, the low Fv/Fm was restored to wild-type levels in the *g-pifi_1* × *pnsl1* double mutant, which was consistent with the recovery of leaf color (Figure 1A). Furthermore, *g-pifi_1* displayed higher Fv/Fm values in lower (older) leaves than in upper (younger) leaves, and showed a negative correlation with NDH activity (Supplemental Figure S4). These data suggest that the decrease in Fv/Fm and the pale-green leaf color in *PIFI*-deficient mutants are attributable to NDH activity.

**Figure 5.**
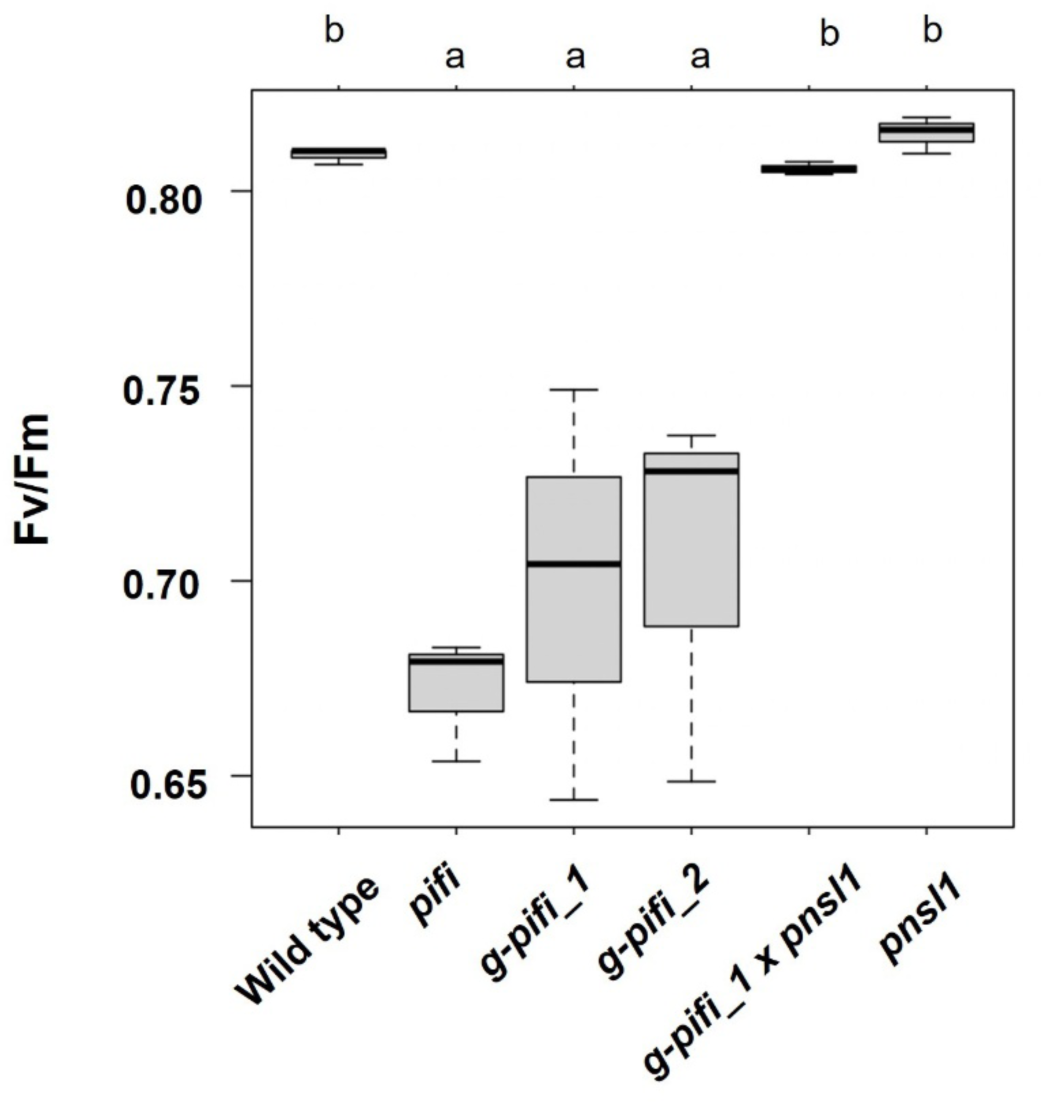
Photoinhibition of PSII activity and accumulation of representative photosynthesis-related proteins in *PIFI*- and *NDH-* deficient mutants. The maximum quantum yield of PSII (Fv/Fm) was measured in attached leaves from each plant. Values are means ± SD (*n* = 3). Statistically significant differences between WT and mutants were determined using the Tukey-Kramer test following one-way ANOVA (*p* < 0.05).

Immunoblot analysis suggested that these decreases in Fv/Fm in *pifi* and *g-pifi* mutants were not due to altered accumulation of photosynthetic proteins (Figure 1B). We further investigated whether photodamage and repair of the D1 protein were affected in *pifi* and *g-pifi* mutants, leading to a decrease in Fv/Fm. PSII activity is maintained by a balance between light-induced damage to the D1 protein and its repair. These processes can be examined separately using lincomycin, which inhibits chloroplast protein synthesis (Nixon et al. 2010; Su et al. 2023). There was no significant difference in the decrease in Fv/Fm between the WT and mutants in the presence or absence of lincomycin, indicating that susceptibility to photodamage and PSII repair were not affected in *pifi* and *g-pifi* mutants (Supplemental Figure S6). Despite a lower Fv/Fm, the quantum yield of PSII (Φ_II_), electron transport rate from PSII (ETR), non-photochemical quenching (NPQ), and PSII reaction center closure (1 - qP) under steady-state conditions in *pifi* and *g-pifi* mutants were similar to those of wild-type (Supplemental Figure S7). These data suggest that PSII function under light illumination is not substantially affected by the lack of PIFI.

We then examined whether PQ reduction in darkness affected PSII efficiency, particularly in the partially disassembled PSII. A previous report has suggested that heat treatment or the absence of extrinsic subunits in PSII allows the reduced PQ pool to drive a back reaction that reduces the Mn cluster in the dark, leading to disassembly and a marked decline in PSII activity (Haveux 1992; Haveux 1996; Ifuku et al. 2005). After dark heat treatment, the Fv/Fm values in the *pifi*, *g-pifi_1*, and *g-pifi_2* mutants were significantly lower than those in the wild type (WT) (Figure 6). Consistent with a previous report (Marutani et al. 2012), this decrease was largely recovered by light illumination and was less pronounced in the *pnsl1* mutant, which lacks NDH, and therefore exhibits less PQ reduction than the WT. The *g-pifi_1* × *pnsl1* double mutant also exhibited a smaller decrease in Fv/Fm. These data suggest that NDH-mediated PQ pool reduction in darkness in *PIFI*-deficient mutants would affect PSII efficiency.

**Figure 6.**
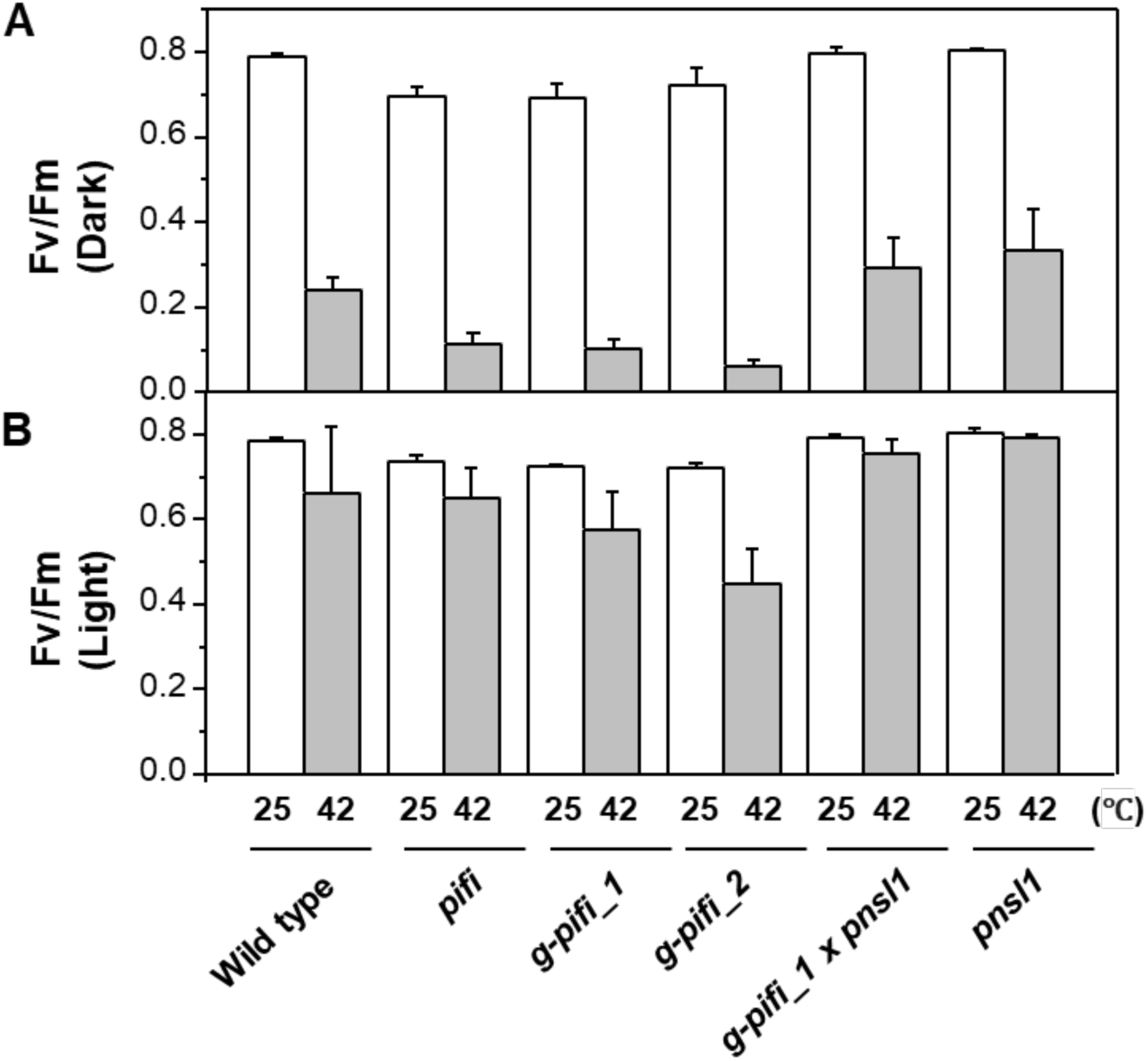
Evaluation of PQ pool reduction using Fv/Fm under light and dark conditions before and after heat stress. Fv/Fm was measured under low light (10 µmol photons m⁻² s⁻¹; A) and in the dark (B) in the wild type and mutant strains. Measurements were performed before heat treatment (25 °C) and after heat stress at 42 °C. Values represent means ± SD (*n* = 3).

### Fv/Fm is not affected in the *trxm4* mutant, showing PQ pool reduction

To confirm that the reduction in the PQ pool by increased NDH activity affects PSII efficiency, we characterized a mutant lacking thioredoxin (Trx) m4 (*trxm4*). It has been reported that in *trxm4*, NDH activity also increases Chl fluorescence after AL is turned off, which is canceled by FR (Courteille et al. 2013). Trxis a disulfide reductase that controls the redox state of several photosynthetic proteins, and TRXm4 is one of its isoforms (Lemair et al. 2007; Schürmann et al. 2008). The increase in Chl fluorescence via NDH activity caused a PQ reduction in the dark in the *trxm4* mutant after AL was turned off, and high Chl fluorescence was maintained, as observed in the *PIFI*-deficient mutants (Figure 7A). Furthermore, the phosphorylation level of LHCII in the dark was higher for *trxm4*, suggesting greater PQ reduction (Figure 7D). However, the Fv/Fm of *trxm4* was similar to that of the WT (Figure 7B). It should be noted that, unlike in *PIFI*-deficient mutants, the NDH-PSI complex was not destabilized in *trxm4* (Figure 7C). These results suggested that the reduction of PQ by NDH alone does not necessarily affect PSII, whereas PQ reduction by the destabilized NDH complex affects PSII efficiency in the absence of PIFI.

**Figure 7.**
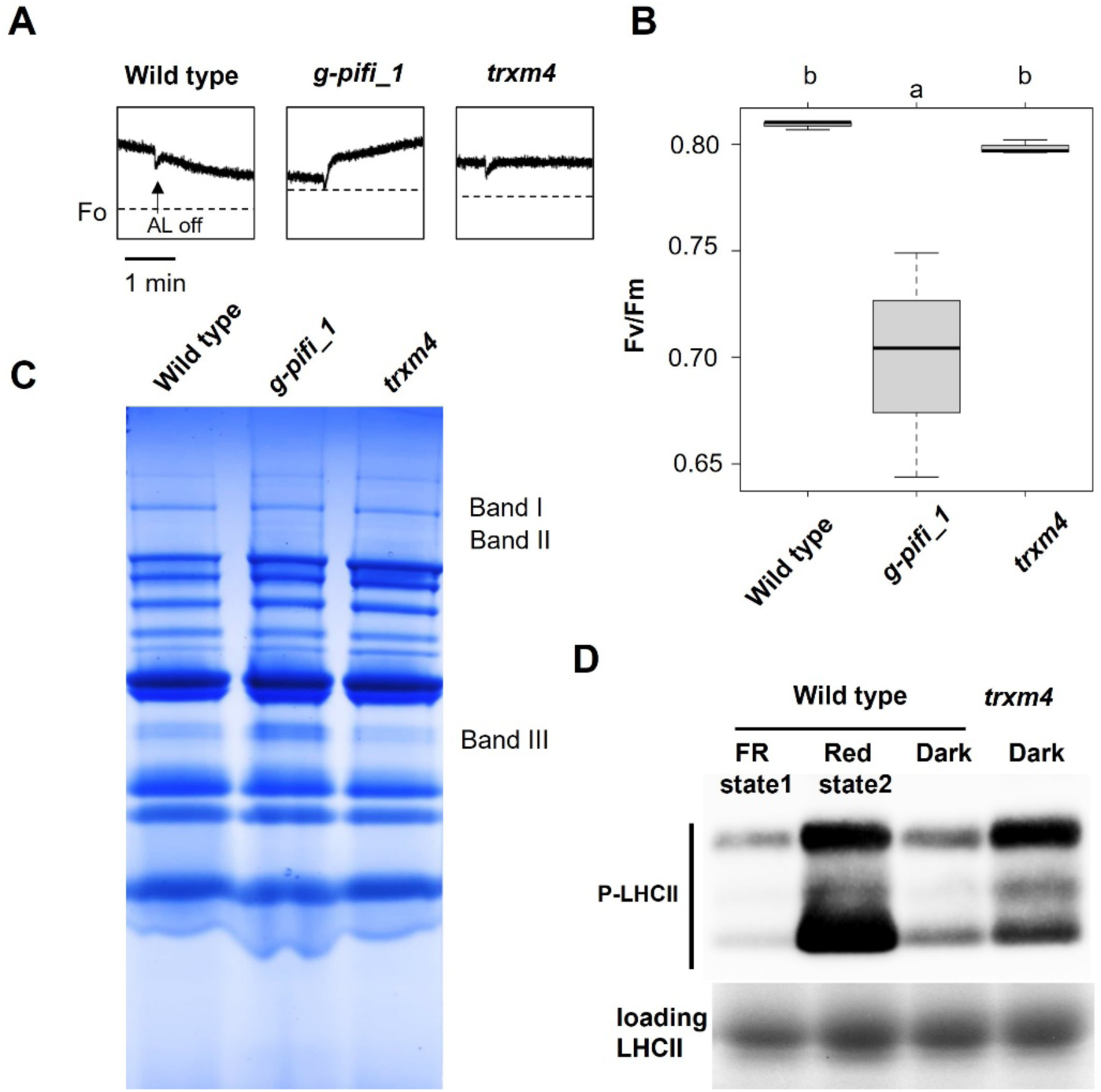
Comparison of phenotypes between *g-pifi_1* and *trxm4*. A. Post-illumination Chl *a* fluorescence used to evaluate NDH activity in WT, *g-pifi_1*, and *trxm4*. B. Fv/Fm was measured in attached leaves from each plant. Values are means ± SD (*n* = 3). Statistically significant differences between WT and mutants were determined using the Tukey-Kramer test following one-way ANOVA (*p* < 0.05). C. Principal photosynthesis protein complexes were separated using BN-PAGE. Thylakoids were isolated from the leaves of wild-type (Col-0) and mutant plants, and then solubilized with 1.0% (w/v) n-dodecyl-β-D-maltoside. Proteins corresponding to 10 µg of Chl were separated on a 3–12% gradient gel. D. Phosphorylation state of LHCII under dark in wild type and *trxm4*. Thylakoid membrane proteins separated using SDS-PAGE and immunoblotted with an anti-phospho-threonine antibody to detect phosphorylated LHCII (P-LHCII). In the wild type, LHCII was dephosphorylated under far-red light (state 1) and phosphorylated under red light (state 2), serving as negative and positive controls, respectively. The lower panel shows total LHCII as a loading control.

## Discussion

In this study, the molecular function of the thylakoid protein PIFI was examined in *Arabidopsis thaliana* using genome-edited mutant strains and the T-DNA insertion strain (Figure 1, Supplemental Figure S1). All *PIFI*-deficient mutants, including the T-DNA insertion strain, exhibited an enhanced post-illumination Chl fluorescence increase, pale-colored leaves, and decreased Fv/Fm of PSII (Figures 1 and 5). In contrast, destabilization of the NDH-PSI supercomplex was evident in *PIFI*-deficient mutants (Figure 3). In the double mutant lacking both PIFI and NDH, NDH activity was absent, and leaf color and Fv/Fm recovered to wild-type levels. These findings suggest that PIFI plays a role in maintaining the stability of the NDH-PSI supercomplex and in regulating the redox state of the PQ pool in darkness, both of which are essential for optimal PSII efficiency.

We showed that PIFI is a thylakoid-localized protein associated with the NDH-PSI supercomplex (Figure 4A, C). PIFI was detected not only in this fraction, but also in the lower molecular weight fractions (Figure 4B), indicating that PIFI can independently accumulate in thylakoids. However, PIFI accumulation was lower in the *pnsl1* mutant lacking the NDH complex (Figure 1B). Therefore, PIFI interacts with the NDH-PSI supercomplex and contributes to its stability, and vice versa. PIFI has not been resolved within the cryo-EM structure of the NDH-PSI supercomplex (Shen et al. 2022; Su et al. 2022; Introini et al. 2025) and exhibits a high turnover rate in chloroplasts (Li et al. 2017). These data suggest that the interaction between PIFI and the NDH-PSI supercomplex can be weak or transient, and that PIFI plays a dynamic rather than a constitutive role in regulating NDH activity.

Paradoxically, NDH activity increased despite the apparent instability of the NDH-PSI supercomplex in *pifi* and *g-pifi* mutants. In typical NDH-deficient mutants such as *crr2* and *pql3*, the lack of an NDH complex abolishes NDH activity, as indicated by the absence of a transient increase in Chl fluorescence in the dark (Munshi et al. 2006; Ishikawa et al. 2020). The *lhca5* and *lhca6* mutants, as well as the *lhca5 lhca6* double mutant, which lack the linker proteins connecting NDH to PSI, retain this transient Chl fluorescence signal in immature leaves but not in mature leaves, indicating that the NDH complex becomes unstable without Lhca5 and Lhca6 (Peng et al. 2009; Peng and Shikanai 2011). In the g-*pifi_1* mutant, PSI dissociated from the NDH complex, while Lhca5 and Lhca6 remained bound to the NDH complex (Figure 3). This leaves the NDH complex detached from PSI in a stable and active form in *PIFI*-deficient mutants (Figure 5), resulting in deregulated NDH activity.

Deregulated NDH activity in *pifi* and *g-pifi* mutants led to elevated LHCII phosphorylation in the dark and a lower Fv/Fm of PSII (Figure 7). In contrast, the *trxm4* mutant also exhibited elevated NDH activity and LHCII phosphorylation but maintained Fv/Fm (Figure 7B). As the NDH-PSI supercomplex is maintained in *trxm4*, PQ reduction by the NDH complex dissociated from PSI may be responsible for the lower Fv/Fm in *PIFI*-deficient mutants. PSI and NDH are localized in the stroma lamellae in the thylakoid membranes of land plants, whereas PSII is concentrated in the grana stacks (Chow et al. 2005). The physiological role of NDH-PSI supercomplex formation has been considered to stabilize NDH and may facilitate CEF-PSI via Fd (Su et al. 2022). Our data further suggest that the association of NDH with PSI is important for controlling PQ reduction by reducing Fd, and that PIFI is involved in this regulation to prevent adverse effects on PSII.

To date, PIFI has only been identified in angiosperms that have complex I-type NDH in chloroplasts. In terrestrial plants, NDH-mediated reduction of the PQ pool should be suppressed in darkness, because maintaining the PQ pool in an oxidized state is essential for proper photosynthetic regulation (Havaux 2020). In contrast, dark reduction of the PQ pool via respiratory electron input has been observed in some cyanobacteria and eukaryotic algae (Aoki and Katoh 1982). Although thylakoids in cyanobacteria and eukaryotic algae lack stacked grana, these organisms have developed specialized membrane compartments for PSII biogenesis and assembly, such as the thylakoid center or translation zone (T-zone) (Nickeksen and Rengstl; 2013, Komenda et al., 2024), where the negative effect of the reduced PQ pool may be escaped. In land plants, occur in stromal lamellae, where. PQ diffusion is thought to be less constrained that in grana stacks but still influenced by membrane microdomains formed by dense protein complexes, including PSII and LHCII (Lavergne et al. 1992; Kirchhoff 2013; Krichhoff et al. 2000). This organization difference requires precise regulation of NDH-dependent PQ reduction in the stromal lamellae to prevent the undesired reduction of immature PSII complexes. Further studies are required to elucidate how PIFI interacts with and regulates the activity of the PSI-NDH complex.

## Material and methods

### Generation of the *g-pifi* mutant using the CRISPR/Cas9-mediated gene knockout method

CRISPR-Cas9-mediated genome editing was used to generate *g-pifi_1* and *g-pifi_2* mutants (Tsutsui and Higashiyama 2017). Two regions of the *PIFI* coding sequence, positions 427–450 and 35–58 relative to the translation initiation site, were selected as the target sites for the single guide RNAs. The complementary oligonucleotides of these target sites were annealed and ligated to the *Aar*I-treated *pKIR1.1*, using the Mighty Mix DNA Ligation Kit (Takara, Shiga, Japan). The following oligonucleotides were used: 5′-ATTGGGTTGGTGGAAGACCATTCA-3′ and 5′-AAACTGAATGGTCTTCCACCAACC-3′ for *g-pifi_1*, and 5′-ATTGGGATAAAGGCTGCGACGGAG-3′ and 5′-AAACCTCCGTCGCAGCCTTTATCC-3′ for *g-pifi_2*. The resulting constructs were introduced into the WT (Col-0) via *Agrobacterium*-mediated transformation using the floral dip method (Clough et al. 1998). We selected T2 plants harboring heterozygous mutations at the target sites from which T_3_ seeds were obtained. Among the T_3_ plants, we identified homozygous mutations. To identify homozygous *g-pifi_1* and *g-pifi_2* mutations, genomic fragments of the *PIFI* gene were amplified using the following oligonucleotides: 5′-TCCTTCTTTCATCCCCAAAA-3′ and 5′-TGGTAGCCCCATTAAACAGC-3′. The resulting PCR products were subjected to DNA sequencing, and T_3_ plants carrying homozygous *pifi* mutations were isolated.

### Plant materials and growth conditions

Arabidopsis (*Arabidopsis thaliana*) ecotype Columbia-0 was used as the wild type and background. Seeds of Arabidopsis with T-DNA-inserted *pifi* (SALK_085656) and *pnsl1* (SALK_0020674) were obtained from the Arabidopsis Biological Resource Center (Wang and Prots 2007; Ishikawa et al. 2007). Two genome-edited *pifi* strains (*g-pifi_1* and *g-pifi_2*) were generated, as described above. To create a double mutant, *g-pifi_1* was crossed with *pnsl1*. The presence of T-DNA insertions was confirmed by PCR (for primers, see Supplementary Table S1). Genome-edited mutations were confirmed using Sanger sequencing (Supplemental Figure S1).

Plants were grown in soil after germinating on Petri dishes containing Murashige and Skoog medium with 1.0% (w/v) agar and 1% (w/v) sucrose and grown for 3 weeks in growth chambers (80 µmol photons m^−2^ s^−1^, 16 h light/8 h dark cycles or continuous light, 22 °C).

### Measurement of NDH activities

For the measurements, four-week-old plants were dark-adapted for 15 min. The NDH activity was measured using a PAM Chl fluorescence analysis system (PAM-2000; MINI-PAMII; Walz, Effeltrich, Germany). Photosynthetically active light (AL, LED lamp) was applied for 4 min, followed by light measurement (ML). The transient increase in Chl fluorescence observed under these conditions was used as an indicator of NDH activity. FR light exposure was conducted 90 seconds after turning off the AL, for 60 seconds. FR light can cancel the PQ pool reduction by specifically exciting PSI. A decrease in Chl fluorescence induced by FR light indicated that the signal depended on PQ pool reduction.

### Chl fluorescence analysis

Four-week-old plants were dark-adapted for over 1 h, and their photosynthetic profiles were measured using a Dual-PAM (Walz, Effeltrich, Germany). Fv/Fm, Φ_II_, ETR, NPQ, and PSII reaction center closure (1 − qP) under steady-state were determined as described previously (Ishihara et al. 2007; Gu et al. 2017).

### Heat treatment for evaluating PQ pool redox state

After 30 min of dark adaptation, leaves were detached from the plants and subjected to heat treatment using a Peltier device (VPE-35; VICS, Japan). The leaves were treated at 42 °C for 30 min, either with or without light, followed by 15 min of dark adaptation at 25 °C before measuring Fv/Fm. For control samples, the leaves were kept at 25 °C for 30 min rather than subjected to a 42 °C heat treatment.

### Whole protein extraction

Leaves were ground in liquid nitrogen and solubilized for SDS-PAGE in 1× SDS sample buffer [62.5 mM Tris–HCl (pH 6.8), 2.5% (w/v) SDS, 10% (w/v) glycerol, 2.5% (v/v) 2-mercaptoethanol, and a trace amount of bromophenol blue (BPB)] at 95 °C for 5 min. Protein content was measured using Bradford reagent (Bio-Rad, Hercules, CA, USA).

### SDS-PAGE and Immunoblot analysis

Whole-leaf protein (20 µg) was loaded in each lane of a 15% polyacrylamide gel and separated using SDS-PAGE. Proteins were transferred onto Immobilon-P PVDF membranes (Millipore, Billerica, MA, USA) and incubated with specific antibodies, as shown in the relevant figure legends. Specific antibodies against PNSL1 and PnsB2 were prepared as described previously (Ishihara et al., 2007; Takabayashi et al., 2009). For D1, Lhcb3, PsaA, and Lhca1, commercially available polyclonal antibodies (Agrisera, Vännäs, Sweden) were used. Phosphorylated LHCII (P-LHCII) was detected using an anti-phospho-threonine antibody (Cell Signaling tech. MA, USA). A specific antibody against PIFI was generated using the full-length PIFI protein as an antigen.

### Chl measurements

Leaves of identical fresh weight were ground in a mortar and pestle using liquid N. Photosynthetic pigments were extracted using 80% acetone. Chl content was determined as previously described (Porra et al. 1989).

### Isolation of intact chloroplasts and subfractionation

Intact chloroplasts were isolated from 3-week-old Col-0 leaves using the two-layer Percoll density gradient method as previously reported (Imaizumi et al. 2025). Chloroplasts were osmotically lysed by incubation in ice-cold hypotonic buffer (50 mM HEPES-KOH, 1 mM MnCl_2_, 1 mM MgCl_2_, and 2 mM EDTA, pH 8.0) for 30 min, and fractionated into stroma and thylakoids by centrifugation. The thylakoids were washed three times with hypotonic buffer to minimize stromal protein contamination.

### Isolation of the thylakoid membrane

Arabidopsis rosette leaves (3 g) were homogenized in 20 mL of ice-cold ground buffer [50 mM HEPES-KOH (pH 7.5), 0.33 M sorbitol, 2 mM EDTA, 1 mM MgCl_2_, 0.05% (w/v) BSA, 5 mM sodium-ascorbate] using a Polytron P10-35GT (Kinematica AG, Malters, Switzerland). The homogenate was filtered through Miracloth (Merck KGaA, Germany) and centrifuged at 2,500 × *g* at 4 °C for 5 min. The pellets were suspended in 5 ml of the Grind buffer and centrifuged at 2,500 × *g* at 4 °C for 5 min. The pellets were resuspended in 5 ml of Lysis Buffer [10 mM HEPES-KOH (pH 8.0)] and centrifuged at 2,500 × *g* at 4 °C for 5 min to obtain the thylakoid membrane fraction. All steps were performed in the dark, and instruments and reagents were cooled on ice.

### BN-PAGE and 2D SDS-PAGE

The isolated thylakoids were thawed on ice, solubilized with 1.0% (w/v) *n*-dodecyl-β-D-maltoside containing 10% (v/v) glycerol. The supernatants were corrected after centrifugation at 2,500 × *g* at 4 °C for 5 min, and the BN-sample loading buffer containing CBB G-250 was added to the sample. The sample corresponding to 10 µg Chl was loaded in each lane of a 3–12% Bis–Tris Gel (Invitrogen, CA, USA), described in (Ishikawa et al., 2020; Takeuchi et al. 2025). After electrophoresis, each lane of the gel was excised and the gel was solubilized in 2× SDS sample buffer [125 mM Tris–HCl (pH 6.8), 5% (w/v) SDS, 20% (w/v) glycerol, 5% (v/v) 2-mercaptoethanol, and a trace amount of BPB] and incubated at 65 °C for 15 min after 37 °C for 15 min with shaking. The solubilized gel was applied to a 12% polyacrylamide gel containing urea. The immunoblot analysis was performed as described above. Thylakoid membranes were extracted on the day of BN-PAGE to minimize the effects of thylakoid membrane decomposition.

### In-gel digestion and mass spectrometry

For MS analysis, gel lanes of bands I, II, and III were cut into gel pieces from the wild-type, *pifi*, and *g-pifi_1.* The gel pieces were reduced and alkylated, followed by digestion with trypsin as previously described. The MS analysis was performed using a Q Exactive Plus Mass Spectrometer (Thermo Scientific, Bremen, Germany). The LC-MS/MS raw files were processed using Mascot software (Matrix Science, U.K.).

### Analysis of PSII photoinhibition

Leaf discs were prepared from the leaves of four-week-old plants and floated in 1 mL of 10 mM sodium phosphate buffer (pH 7.0, NaOH) containing 1 mM lincomycin, an inhibitor of translation in plastids. The inhibitor was then infiltrated into the leaf disks using a vacuum desiccator for 5 min. DMSO was added to control samples without lincomycin. The samples were subjected to high-light treatment at an intensity of 550 µmol photons m^−2^s^−1^. The Fv/Fm ratio was measured using Mini-PAM (Walz, Germany) after 30, 60, 120, and 180 min of light treatment.

### Accession numbers

Sequence data from this study can be found in the National Center for Biotechnology Information (NCBI) database under the accession numbers PIFI (At3g15840).

## Acknowledgments

We thank Prof. Ken Motohashi (Kyoto Sangyo University) for kindly providing *trxm4* mutant seeds. We also thank Mr. Yuzo Watanabe (Kyoto University) for performing the mass spectrometry analysis.

## Author contributions

Ke.I., K.Y., and T.H. conceived the project. M.M., Ko.I., K.M., A.K, Y.C., N.I., and K.K performed the experiments. M.M., Ko.I. and K.K. analyzed the data. K.K. and Ke.I wrote the manuscript with input from all authors. All authors read and approved the manuscript.

## Supplementary data

The following materials are available in the online version of this article.

**Supplementary Figure S1.** Mutation sites in T-DNA inserted mutant *pifi* and genome-edited mutants *g-pifi_1* and *g-pifi_2*.

**Supplementary Figure S2.** Comparison of Chl content of wild type, T-DNA inserted *pifi*, two strains of genome-edited *pifi* (*g-pifi_1*, *g-pifi_2*), and *g-pifi_1* crossed with *pnsl1*, an NDH-deficient mutant.

**Supplementary Figure S3.** Repeats of NDH activity. Post-illumination Chl *a* fluorescence was used to evaluate NDH activity.

**Supplementary Figure S4.** Comparison of NDH activity among leaves of different ages within a single plant.

**Supplementary Figure S5.** Comparative analysis of the formation of the NDH-PSI supercomplex in wild-type and *PIFI*-deficient mutant (*g-pifi_1*).

**Supplementary Figure S6.** Evaluation of Fv/Fm using lincomycin, a PSII repair inhibitor.

**Supplementary Figure S7.** Light-dependent photosynthetic responses in the wild type and *pifi*-related mutants.

**Supplementary Table S1.** Number of peptide subunits of PSI, LHCA, and NDH in each band.

## REFERENCES

Alric J, Johnson X. Alternative electron transport pathways in photosynthesis: a confluence of regulation. Curr Opin Plant Biol. 2017:37: 8–86.

Aoki M, Katoh S. Oxidation and reduction of plastoquinone by photosynthetic and respiratory electron transport in a cyanobacterium *Synechococcus* sp. Biochimica et Biophysica Acta (BBA) - Bioenergetics 1982:682:307–314.

Asada K. THE WATER-WATER CYCLE IN CHLOROPLASTS: Scavenging of Active Oxygens and Dissipation of Excess Photons. Annu Rev Plant Physiol Plant Mol Biol. 1999:50:601–639.

Bennoun P. Evidence for a respiratory chain in the chloroplast. Proc Natl Acad Sci U S A. 1982: 79(14):4352–4356.

Ben-Shem A, Frolow F, Nelson N. Crystal structure of plant photosystem I. Nature. 2003: 426(6967):630–635.

Burrows PA, Sazanov LA, Svab Z, Maliga P, Nixon PJ. Identification of a functional respiratory complex in chloroplasts through analysis of tobacco mutants containing disrupted plastid ndh genes. EMBO J. 1998:17:868–876.

Chow WS, Kim EH, Horton P, Anderson JM. Granal stacking of thylakoid membranes in higher plant chloroplasts: the physicochemical forces at work and the functional consequences that ensue. Photochem Photobiol Sci. 2005:4(12):1081–1090.

Clough SJ, Bent AF. Floral dip: a simplified method for Agrobacterium-mediated transformation of Arabidopsis thaliana. Plant J. 1998:16(6):735–743.

Courteille A, Vesa S, Sanz-Barrio R, Cazalé AC, Becuwe-Linka N, Farran I, Havaux M, Rey P, Rumeau D. Thioredoxin m4 controls photosynthetic alternative electron pathways in Arabidopsis. Plant Physiol. 2013:161(1):508–520.

Dal Corso G, Pesaresi P, Masiero S, Aseeva E, Schünemann D, Finazzi G, Joliot P, Barbato R, Leister D. A complex containing PGRL1 and PGR5 is involved in the switch between linear and cyclic electron flow in Arabidopsis. Cell. 2008:132:273–285.

Degen GE, Jackson PJ, Proctor MS, Zoulias N, Casson SA, Johnson MP. High cyclic electron transfer via the PGR5 pathway in the absence of photosynthetic control. Plant Physiol. 2023:192(1):370–386.

Ermakova M, Woodford R, Fitzpatrick D, Nix SJ, Zwahlen SM, Farquhar GD, von Caemmerer S, Furbank RT. Chloroplast NADH dehydrogenase-like complex-mediated cyclic electron flow is the main electron transport route in C_4_ bundle sheath cells. New Phytol. 2024:243(6):2187–2200.

Gu J, Zhou Z, Li Z, Chen Y, Wang Z, Zhang H, Yang J. Photosynthetic Properties and Potentials for Improvement of Photosynthesis in Pale Green Leaf Rice under High Light Conditions. Front Plant Sci. 2017:8:1082.

Hashimoto M, Endo T, Peltier G, Tasaka M, Shikanai T. A nucleus-encoded factor, CRR2, is essential for the expression of chloroplast ndhB in Arabidopsis. Plant J. 2003:36(4):541–549.

Havaux M. Plastoquinone in and beyond photosynthesis. Trends Plant Sci. 2020:25(12):1252–1265.

Hertle AP, Blunder T, Wunder T, Pesaresi P, Pribil M, Armbruster U, Leister D. PGRL1 is the elusive ferredoxin-plastoquinone reductase in photosynthetic cyclic electron flow. Mol Cell. 2013:49:511–523.

Ifuku K, Yamamoto Y, Ono TA, Ishihara S, Sato F. PsbP protein, but not PsbQ protein, is essential for the regulation and stabilization of photosystem II in higher plants. Plant Physiol. 2005:139(3):1175–1184.

Ifuku K, Endo T, Shikanai T, Aro EM. Structure of the chloroplast NADH dehydrogenase-like complex: nomenclature for nuclear-encoded subunits. Plant Cell Physiol. 2011:52(9):1560–1568.

Imaizumi K, Takagi D, Ifuku K (2025) Antimycin A induces light hypersensitivity of PSII in the presence of quinone Q_B_-site binding herbicides. Plant Physiol.197:kiaf082.

Introini B, Hahn A, Kühlbrandt W. Cryo-EM structure of the NDH-PSI-LHCI supercomplex from Spinacia oleracea. Nat Struct Mol Biol. 2025:2(6):968–978.

Ishihara S, Takabayashi A, Ido K, Endo T, Ifuku K, Sato F. Distinct functions for the two PsbP-like proteins PPL1 and PNSL1 in the chloroplast thylakoid lumen of Arabidopsis. Plant Physiol. 2007:145(3):668–679.

Ishikawa N, Takabayashi A, Noguchi K, Tazoe Y, Yamamoto H, von Caemmerer S, Sato F, Endo T. NDH-Mediated Cyclic Electron Flow Around Photosystem I is Crucial for C_4_ Photosynthesis. Plant Cell Physiol. 2016:57(10):2020–2028.

Ishikawa N, Yokoe Y, Nishimura T, Nakano T, Ifuku K. PsbQ-Like Protein 3 Functions as an Assembly Factor for the Chloroplast NADH Dehydrogenase-Like Complex in Arabidopsis. Plant Cell Physiol. 2020:61(7):1252–1261.

Joët T, Genty B, Josse EM, Kuntz M, Cournac L, Peltier G. Involvement of a plastid terminal oxidase in plastoquinone oxidation as evidenced by expression of the Arabidopsis thaliana enzyme in tobacco. J Biol Chem. 2002:277(35):31623–31630.

Johnson GN. Physiology of PSI cyclic electron transport in higher plants. Biochim Biophys Acta 2011:1807:384–389.

Johnson MP. Structure, regulation and assembly of the photosynthetic electron transport chain. Nat. Rev. Mol. Cell Biol. 2025:26(9):667–690.

Kato Y, Odahara M, Fukao Y, Shikanai T. Stepwise evolution of supercomplex formation with photosystem I is required for stabilization of chloroplast NADH dehydrogenase-like complex: Lhca5-dependent supercomplex formation in *Physcomitrella patens*. Plant J. 2018:96(5):937–948.

Kato Y, Odahara M, Shikanai T. Evolution of an assembly factor-based subunit contributed to a novel NDH-PSI supercomplex formation in chloroplasts. Nat Commun. 2021:12(1):3685.

Kono M, Terashima I. Long-term and short-term responses of the photosynthetic electron transport to fluctuating light. J Photochem Photobiol B. 2014:137:89–99.

Kirchhoff H., Horstmann S., Weis E. Control of the photosynthetic electron transport by PQ diffusion microdomains in thylakoids of higher plants. Biochim Biophys Acta. 2000:1459: 148–168.

Kirchhoff H. Architectural switches in plant thylakoid membranes. Photosynth Res. 2013:16:481–487.

Kohzuma K, Muraoka S, Kumazawa M, Ifuku K. Evolution and regulatory diversification of plastid F_1_F_O_-ATP synthase. Plant Cell Physiol. 2025:66(11):1619–1632.

Komenda J, Sobotka R, Nixon PJ. The biogenesis and maintenance of PSII: Recent advances and current challenges. Plant Cell. 2024:36(10):3997–4013.

Konno H, Murakami-Fuse T, Fujii F, Koyama F, Ueoka-Nakanishi H, Pack CG, Kinjo M, Hisabori T. The regulatory cysteine pair of the γ subunit is located near the rotor of the chloroplast ATP synthase. J Biol Chem. 2006:281:35236–35241.

Kouřil R, Strouhal O, Nosek L, Lenobel R, Chamrád I, Boekema EJ, Šebela M, Ilík P. Structural characterization of a plant photosystem I and NAD(P)H dehydrogenase supercomplex. Plant J. 2014:77(4):568–576.

Krieger-Liszkay A, Shimakawa G. Regulation of the generation of reactive oxygen species during photosynthetic electron transport. Biochem Soc Trans. 2022:50(2):1025–1034.

Krysiak M, Węgrzyn A, Kowalewska Ł, Kulik A, Ostaszewska-Bugajska M, Mazur J, Garstka M, Mazur R. Light-independent pathway of STN7 kinase activation under low temperature stress in runner bean (Phaseolus coccineus L.). BMC Plant Biol. 2024:4(1):513.

Lavergne D, Nato A, Dupuis JM, Péan M, Chagvardieff P. Evidence for the expression of morphological and biochemical characteristics of C3-photosynthesis in chlorohyllous callus cultures of Zea mays. Physiol. Plant. 1992:84:292–300.

Lemaire SD, Michelet L, Zaffagnini M, Massot V, Issakidis-Bourguet E. Thioredoxins in chloroplasts. Curr Genet. 2007:51(6):343–365.

Li L, Nelson CJ, Trösch J, Castleden I, Huang S, Millar AH. Protein Degradation Rate in *Arabidopsis thaliana* Leaf Growth and Development. Plant Cell. 2017:29(2):207–228.

Marutani Y, Yamauchi Y, Kimura Y, Mizutani M, Sugimoto Y. Damage to photosystem II due to heat stress without light-driven electron flow: involvement of enhanced introduction of reducing power into thylakoid membranes. Planta. 2012:236(2):753–761.

Messant M, Hani U, Lai TL, Wilson A, Shimakawa G, Krieger-Liszkay A. Plastid terminal oxidase (PTOX) protects photosystem I and not photosystem II against photoinhibition in *Arabidopsis thaliana* and *Marchantia polymorpha*. Plant J. 2024:117(3):669–678.

Miyake C, Asada K. Thylakoid-Bound Ascorbate Peroxidase in Spinach Chloroplasts and Photoreduction of Its Primary Oxidation Product Monodehydroascorbate Radicals in Thylakoids, Plant Cell Physiol. 1992:33(5):541–553.

Munekage Y, Hojo M, Meurer J, Endo T, Tasaka M, Shikanai T. PGR5 is involved in cyclic electron flow around photosystem I and is essential for photoprotection in Arabidopsis. Cell. 2002:110:361–371.

Munekage Y, Hashimoto M, Miyake C, Tomizawa K, Endo T, Tasaka M, Shikanai T. Cyclic electron flow around photosystem I is essential for photosynthesis. Nature. 2004:429(6991):579–582.

Munshi MK, Kobayashi Y, Shikanai T. Chlororespiratory reduction 6 is a novel factor required for accumulation of the chloroplast NAD(P)H dehydrogenase complex in Arabidopsis. Plant Physiol. 2006:141(2):737–744.

Nellaepalli S, Kodru S, Tirupathi M, Subramanyam R. Anaerobiosis induced state transition: a non photochemical reduction of PQ pool mediated by NDH in Arabidopsis thaliana. PLoS One. 2012:7(11):e49839.

Nickelsen J, Rengstl B. Photosystem II assembly: from cyanobacteria to plants. Annu Rev Plant Biol. 2013:64:609–635.

Nixon PJ, Michoux F, Yu J, Boehm M, Komenda J. Recent advances in understanding the assembly and repair of photosystem II. Ann Bot. 2010:106(1):1–16.

Ogawa T, Kobayashi K, Taniguchi YY, Shikanai T, Nakamura N, Yokota A, Munekage YN. Two cyclic electron flows around photosystem I differentially participate in C_4_ photosynthesis. Plant Physiol. 2023:191(4):2288–2300.

Otani T, Kato Y, Shikanai T. Specific substitutions of light-harvesting complex I proteins associated with photosystem I are required for supercomplex formation with chloroplast NADH dehydrogenase-like complex. Plant J. 2018:94(1):122–130.

Pan X, Ma J, Su X, Cao P, Chang W, Liu Z, Zhang X, Li M. Structure of the maize photosystem I supercomplex with light-harvesting complexes I and II. Science. 2018:360(6393):1109–1113.

Peng L, Fukao Y, Fujiwara M, Takami T, Shikanai T. Efficient operation of NAD(P)H dehydrogenase requires supercomplex formation with photosystem I via minor LHCI in Arabidopsis. Plant Cell. 2009:21(11):3623–3640.

Peng L, Shikanai T. Supercomplex formation with photosystem I is required for the stabilization of the chloroplast NADH dehydrogenase-like complex in Arabidopsis. Plant Physiol. 2011:155(4):1629–1639.

Porra RJ, Thompson WA, Kriedemann PE. Determination of accurate extinction coefficients and simultaneous equations for assaying chlorophylls a and b extracted with four different solvents: verification of the concentration of chlorophyll standards by atomic absorption spectroscopy. Biochimica et Biophysica Acta (BBA) – Bioenergetics. 1989:975(3):384–394.

Rog I, Chaturvedi AK, Tiwari V, Danon A. Low light-regulated intramolecular disulfide fine-tunes the role of PTOX in Arabidopsis. Plant J. 2022:109(3):585–597.

Rühle T, Dann M, Reiter B, Schünemann D, Naranjo B, Penzler JF, Kleine T, Leister D. PGRL2 triggers degradation of PGR5 in the absence of PGRL1. Nat Commun. 2021:12(1):3941.

Satoh H, Ohara Y, Hanke G, Ifuku K, Shimakawa G, Suzuki Y, Makino A, Morigaki K, Miyake C. The regulation of PSI cyclic electron transport by both plastoquinone and ferredoxin redox states: correlation with the rate of proton motive force utilization. Front Plant Sci. 2025:16:1626163.

Schürmann P, Buchanan BB. The ferredoxin/thioredoxin system of oxygenic photosynthesis. Antioxid Redox Signal. 2008:10(7):1235–1274.

Shen L, Tang K, Wang W, Wang C, Wu H, Mao Z, An S, Chang S, Kuang T, Shen JR, Han G, Zhang X. Architecture of the chloroplast PSI-NDH supercomplex in *Hordeum vulgare*. Nature. 2022:601(7894):649–654.

Shikanai T, Endo T, Hashimoto T, Yamada Y, Asada K, Yokota A. Directed disruption of the tobacco *ndhB* gene impairs cyclic electron flow around photosystem I. Proc. Natl. Acad. Sci. U. S. A. 1998:95:9705–9709

Shikanai T. Chloroplast NDH: A different enzyme with a structure similar to that of respiratory NADH dehydrogenase. Biochim Biophys Acta. 2016:1857(7):1015–1022.

Shikanai T, Yamamoto H. Contribution of cyclic and pseudo-cyclic electron transport to the formation of proton motive force in chloroplasts. Mol. Plant 2017:10:20–29.

Shikanai T, Ieda H, Kobayashi Y, Tamura MN. The chloroplast NADH dehydrogenase-like complex: evolutionary considerations. Plant Cell Physiol. 2025:66(11):1525–1535.

Sonoike K. Photoinhibition and protection of photosystem I. Plant Cell Physiol. 2025:66(11):1562–1574.

Su J, Jiao Q, Jia T, Hu X. The photosystem-II repair cycle: updates and open questions. Planta. 2023:259(1):20.

Su X, Cao D, Pan X, Shi L, Liu Z, Dall’Osto L, Bassi R, Zhang X, Li M. Supramolecular assembly of chloroplast NADH dehydrogenase-like complex with photosystem I from Arabidopsis thaliana. Mol Plant. 2022:15(3):454–467.

Suorsa M. Cyclic electron flow provides acclimatory plasticity for the photosynthetic machinery under various environmental conditions and developmental stages. Front. Plant Sci. 2015:6:800.

Takabayashi A, Kishine M, Asada K, Endo T, Sato F. Differential use of two cyclic electron flows around photosystem I for driving CO_2_ -concentration mechanism in C_4_ photosynthesis. Proc. Natl. Acad. Sci. U.S.A. 2005:102:16898–16903.

Takeuchi K, Harimoto S, Che Y, Kumazawa M, Satoh H, Maekawa S, Miyake C, Ifuku K. The protective role of chloroplast NADH dehydrogenase-like complex (NDH) against PSI photoinhibition under chilling stress. New Phytol. 2025:248(5):2262–2279.

Tikhonov AN. Induction events and short-term regulation of electron transport in chloroplasts: an overview. Photosynth Res. 2015:125:65–94.

Tikkanen M, Aro EM. Interacting short-term regulatory mechanisms enable the conversion of light energy to chemical energy in photosynthesis. J Exp Bot. 2026:77(4):895–909.

Tsutsui H, Higashiyama T. pKAMA-ITACHI Vectors for Highly Efficient CRISPR/Cas9-Mediated Gene Knockout in Arabidopsis thaliana. Plant Cell Physiol. 2017:58(1):46–56.

Wang D, and Portis Jr. A.R. A novel nucleus-encoded chloroplast protein, PIFI, is involved in NAD(P)H dehydrogenase complex-mediated chlororespiratory electron transport in Arabidopsis. Plant Physiol. 2007:144:1742–1752.

Wang P, Xue L, Batelli G, Lee S, Hou YJ, Van Oosten MJ, Zhang H, Tao WA, Zhu JK. Quantitative phosphoproteomics identifies SnRK2 protein kinase substrates and reveals the effectors of abscisic acid action. Proc Natl Acad Sci U S A. 2013:110(27):11205–11210.

Witt HT. Energy conversion in the functional membrane of photosynthesis. Analysis by light pulse and electric pulse methods. The central role of the electric field. Biochim Biophys Acta. 1979:505(3-4): 55–427.

Yadav KN, Semchonok DA, Nosek L, Kouřil R, Fucile G, Boekema EJ, and Eichacker LA. Supercomplexes of plant photosystem I with cytochrome *b*_6_*f*, light-harvesting complex II and NDH. Biochim Biophys Acta Bioenerg. 2017:1858(1):12–20.

Yamori W, Shikanai T. Physiological functions of cyclic electron transport around Photosystem I in sustaining photosynthesis and plant growth. Annual Review of Plant Biology. 2016:67:81–106.

Yamori W, Makino A, Shikanai T. A physiological role of cyclic electron transport around photosystem I in sustaining photosynthesis under fluctuating light in rice. Sci. Rep. 2016:6:20147.

Zhang S, Zou B, Cao P, Su X, Xie F, Pan X, Li M. Structural insights into photosynthetic cyclic electron transport. Mol Plant. 2023:16(1):187–205.

