## Supplemental for "PIFI Stabilizes Chloroplast NDH–PSI Supercomplex to Maintain Plastoquinone Redox Balance and PSII Efficiency"

**Supplemental Figure S1**

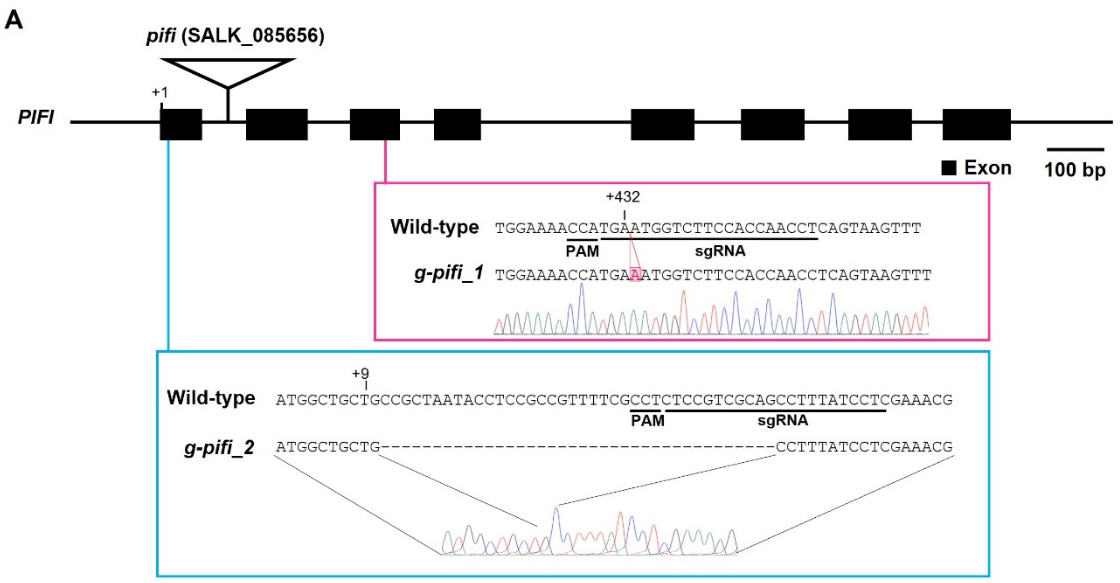


B

```
Wild-type MAAAANTSAV FASPSQPLSS KRSFLYSSRI GPILRRFPRK KLDLQVKAVA TTLAPLEEIK EYKLPSWAMF EMGTAPVYWK
g-pifi_1  MAAAANTSAV FASPSQPLSS KRSFLYSSRI GPILRRFPRK KLDLQVKAVA TTLAPLEEIK EYKLPSWAMF EMGTAPVYWK
g-pifi_2  MAAAFILET* ---------- ---------- ---------- ---------- ---------- ---------- ----------

Wild-type TMNGLPPTSG EKLKLFYNPA ASKLTLNEDY GVAFNGGFNQ PIMCGGEPRA MLKKDRGKAD SPIYTMQICI PKHAVNLIFS
g-pifi_1  TMKWSSTNLR *--------- ---------- ---------- ---------- ---------- ---------- ----------
g-pifi_2  ---------- ---------- ---------- ---------- ---------- ---------- ---------- ----------

Wild-type FTNGVDWDGP YRLQFQVPKR WQNKPIEFFN EGLANELSQD GACERAIFPD SNVVPTRCTM IANLTVEGGD RCNLDLVPGC
g-pifi_1  ---------- ---------- ---------- ---------- ---------- ---------- ---------- ----------
g-pifi_2  ---------- ---------- ---------- ---------- ---------- ---------- ---------- ----------

Wild-type MDTNSEHFNP YANVDDGSCP LELSDSDE*
g-pifi_1  ---------- ---------- ----------
g-pifi_2  ---------- ---------- ----------
```

**Supplementary Figure S1.** Mutation sites in T-DNA inserted mutant *pifi* and genome-edited mutants *g-pifi_1* and *g-pifi_2*. A. Structure of the *PIFI* gene (At3g15840), with exons shown as black boxes and introns as lines. The bar indicates 100 bp. The SALK_085656 strain has a T-DNA insertion in an intron. The genome-edited mutants *g-pifi_1* and *g-pifi_2* were generated using CRISPR-Cas9. The *g-pifi_1* has a single-base insertion at position +432 in the *PIFI* genomic sequence. The *g-pifi_2* has a 38-base deletion at position +9 of the PIFI genomic sequence. The lines on the sequences indicate PAM sequences and sgRNA sequences, respectively. B. Amino acid sequences of the *PIFI* gene in wild type, *g-pifi_1*, and *g-pifi_2*. Asterisks indicate stop codons.

**Supplemental Figure S2**

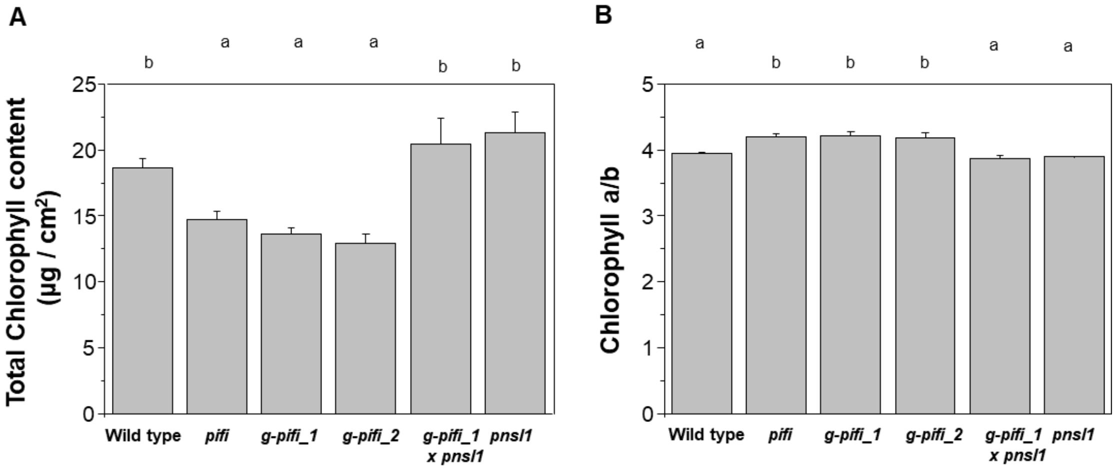


**Supplementary Figure S2.** Comparison of Chl content of wild type, T-DNA inserted *pifi*, two strains of genome-edited *pifi* (*g-pifi_1*, *g-pifi_2*), and *g-pifi_1* crossed with *pnsl1*, an NDH-deficient mutant. A. Total Chl content, and B. Chl a/b ratio. Tukey multiple comparisons test was used to determine significant differences among the plants tested ($P < 0.05$). Bars indicate SD ($n = 3$).

**Supplemental Figure S3**

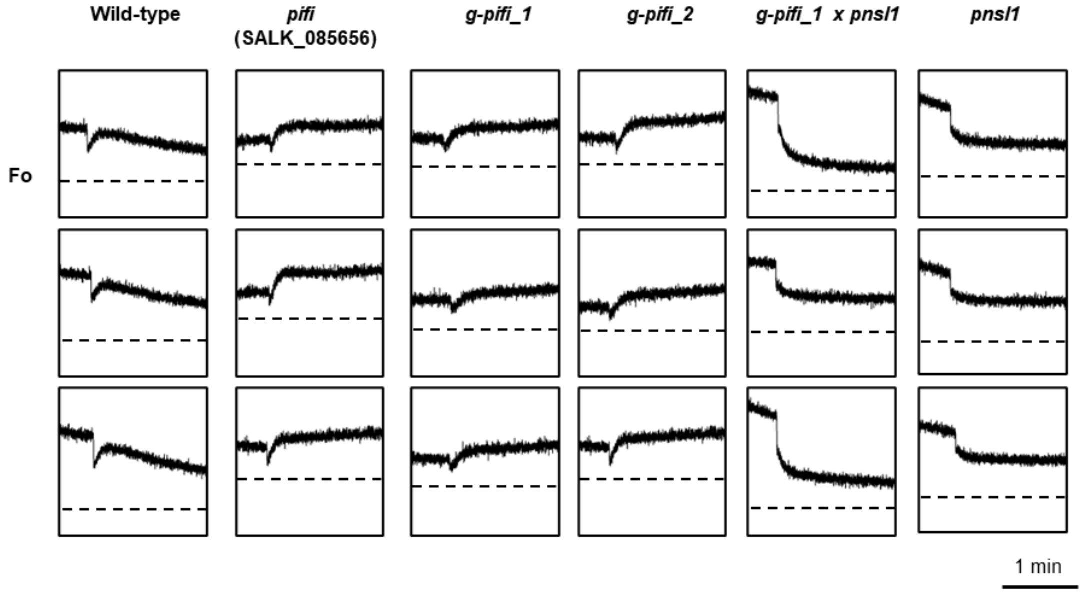


**Supplementary Figure S3.** Repeats of NDH activity. Post-illumination Chl *a* fluorescence used to evaluate NDH activity. The trace represents 1.5 minutes after turning off the actinic light (AL). The dotted line indicates Fo. Bars = 1 min.

**Supplemental Figure S4**

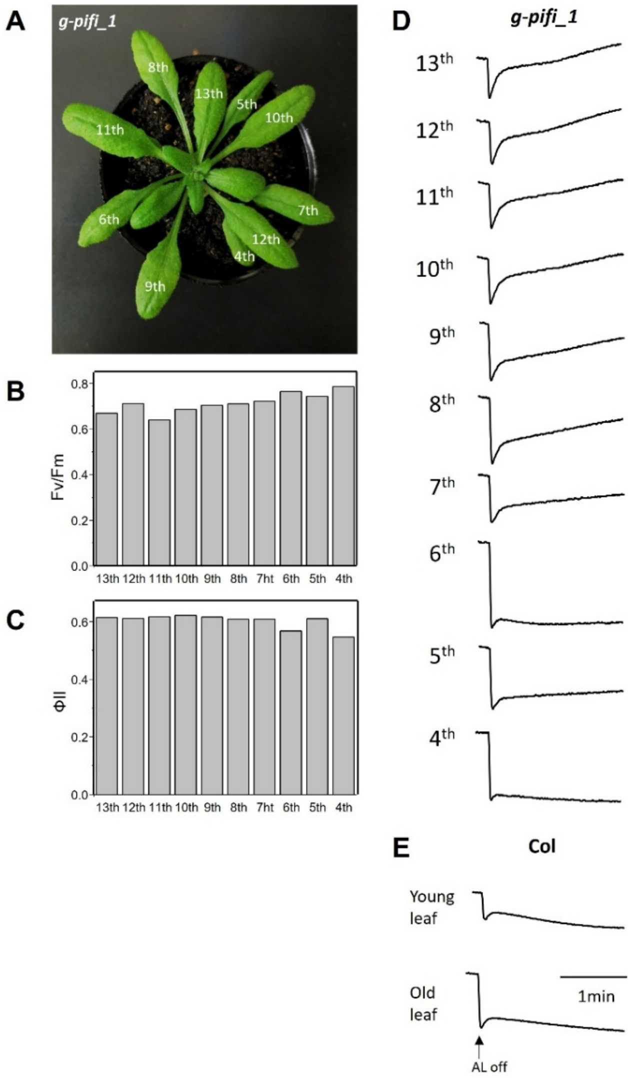


**Supplementary Figure S4.** Comparison of NDH activity among leaves of different ages within a single plant. A. Rosette image of a *g-pifi_1* plant showing leaf positions numbered from the oldest (4th) to the youngest (13th). B. Maximum quantum yield of PSII (Fv/Fm) measured in each leaf. C. Effective quantum yield of PSII ($\Phi_{II}$) measured under actinic light. D. Post-illumination Chl *a* fluorescence traces used to evaluate NDH activity in individual leaves of the *g-pifi_1*. Fluorescence transients vary with leaf age. Traces show fluorescence changes for 2 minutes following the termination of actinic light (AL). E. Representative post-illumination fluorescence kinetics in young and old leaves of the wild-type Col-0 plant. Scale bar = 1 min.

**Supplemental Figure S5**

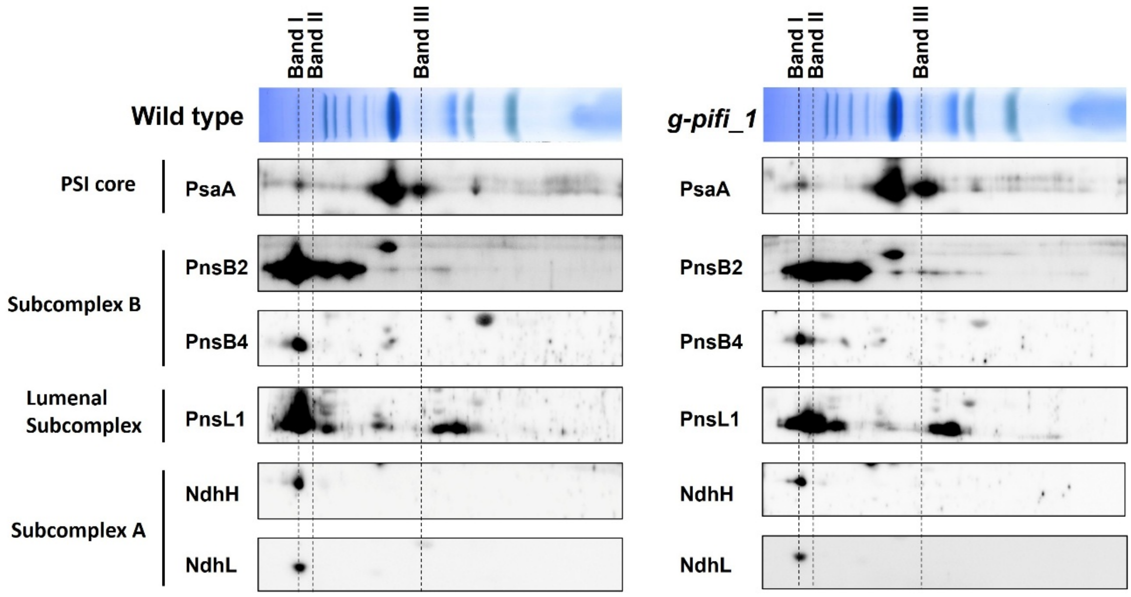


**Supplementary Figure S5.** Comparative analysis of the formation of the NDH-PSI supercomplex in wild-type and *PIFI*-deficient mutant (*g-pifi_1*). Protein complexes separated using BN-PAGE were subsequently subjected to 2D SDS-PAGE in wild-type and *g-pifi_1*. NDH subunits (PnsB2, PnsB4, PnsL1, NdhH, and NdhL) and PSI core (PsaA) were detected using specific antibodies raised against each protein.

**Supplemental Figure S6**

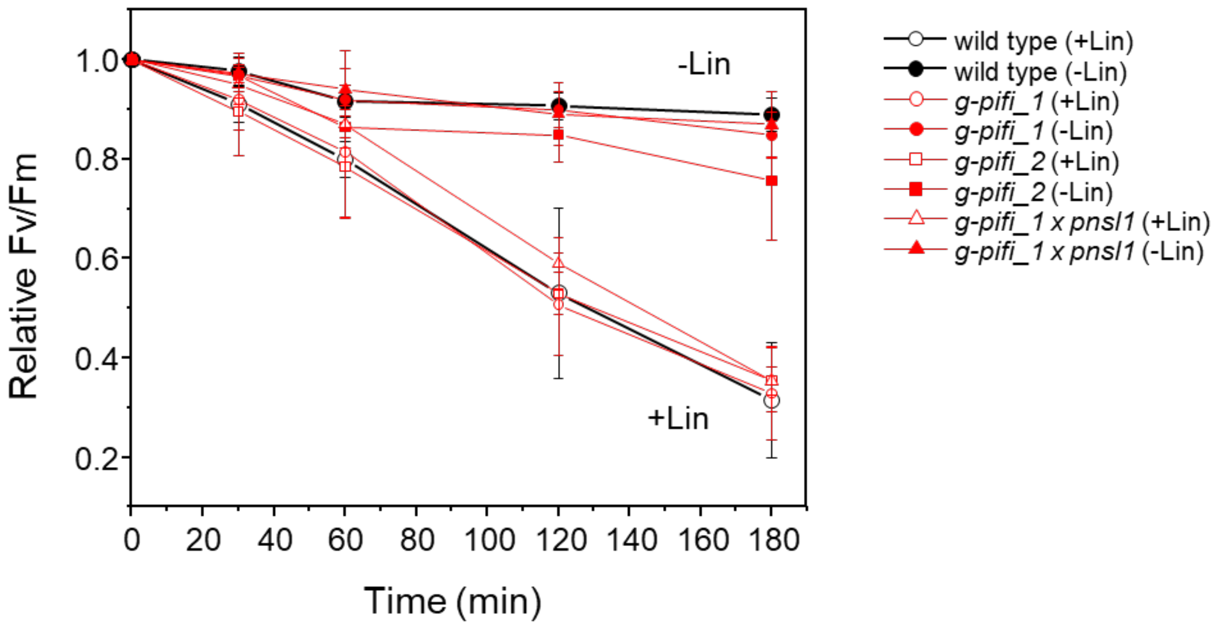


**Supplementary Figure S6**. Evaluation of Fv/Fm using lincomycin, a PSII repair inhibitor. Leaf discs were prepared from leaves of wild type and each mutant, and the time-course changes in the maximum quantum yield of photosystem II, represented by Fv/Fm values, were measured under high light conditions of 550 μmol photons $m^{-2}s^{-1}$. Black symbols and lines are wild type. Red symbols and lines are *PIFI*-deficient mutants. The open symbols indicate values in the absence of lincomycin, while the closed symbols indicate values in the presence of 1 mM lincomycin. Error bars represent SD (*n* = 3).

**Supplemental Figure S7**

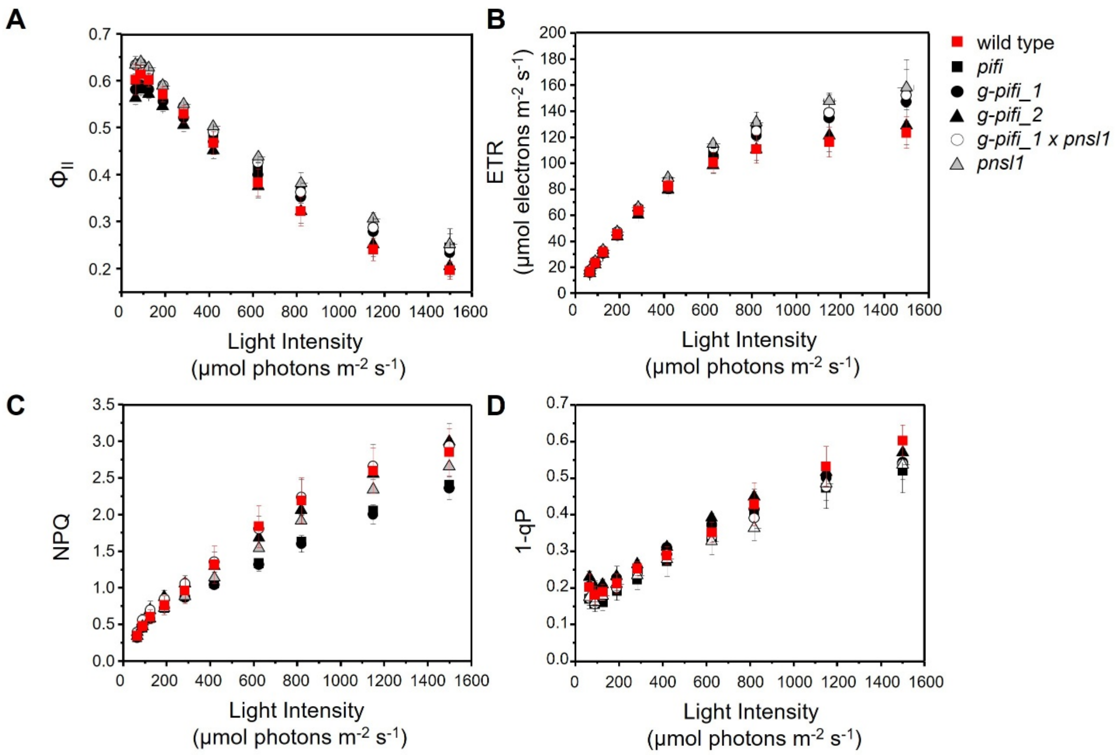


**Supplementary Figure S7.** Light-dependent photosynthetic responses in the wild type and *pifi*-related mutants. A. Effective quantum yield of PSII ($\Phi_{II}$), B. electron transport rate from PSII (ETR), C. non-photochemical quenching (NPQ), and D. fraction of reduced PSII reaction centers (1 − qP) were measured across a range of actinic light intensities (μmol photons $m^{-2}$ $s^{-1}$). Measurements at each light intensity were performed under steady-state conditions after 10 min of illumination. Red squares represent the wild type, black squares represent *pifi*, black circles represent *g-pifi_1*, black triangles represent *g-pifi_2*, open circles represent *g-pifi_1* × *pnsl1*, and open triangles represent *pnsl1*. Data points indicate mean values, and error bars represent ± SE ($n$ = 3).

**Supplemental Table S1**. **Subunit identification by mass spectrometry and Mascot.** Band numbers corresponding to those in Figure 5.

| Band | Assigned subunit | UniProt accession | Protein Mass | Protein Score | | | Protein Matches | | | emPAI | | |
|---|---|---|---|---|---|---|---|---|---|---|---|---|
| | | | | WT | *g-pifi_1* | *g-pifi_1 x pnsl1* | WT | *g-pifi_1* | *g-pifi_1 x pnsl1* | WT | *g-pifi_1* | *g-pifi_1 x pnsl1* |
| **Band I** | **PSI** | | | | | | | | | | | |
| | psaA | P56766 | 83178 | 976 | 907 | 763 | 42 | 34 | 22 | 1.23 | 1.07 | 0.93 |
| | psaB | P56767 | 82423 | 1201 | 1168 | 846 | 47 | 35 | 19 | 1.09 | 1.01 | 0.67 |
| | **Lhca** | | | | | | | | | | | |
| | lhca1 | A8MS75 | 23247 | 275 | 224 | 182 | 11 | 7 | 4 | 0.89 | 0.67 | 0.47 |
| | lhca2 | Q9SYW8 | 27737 | 146 | 59 | 104 | 3 | 2 | 3 | 0.24 | 0.11 | 0.24 |
| | lhca3 | Q9SY97 | 29163 | 438 | 287 | 262 | 11 | 7 | 6 | 1.27 | 0.67 | 0.67 |
| | lhca4 | P27521 | 27716 | 409 | 343 | 332 | 12 | 9 | 7 | 1.12 | 1.12 | 1.12 |
| | lhca5 | Q9C639 | 27784 | 592 | 443 | 0 | 14 | 11 | 0 | 2.63 | 1.93 | 0 |
| | lhca6 | Q8LCQ4 | 29920 | 209 | 154 | 0 | 5 | 3 | 0 | 0.65 | 0.35 | 0 |
| | **NDH** | | | | | | | | | | | |
| | ndhF | P56752 | 85183 | 895 | 714 | 0 | 23 | 17 | 0 | 0.83 | 0.59 | 0 |
| | ndhH | P56753 | 45473 | 1232 | 949 | 0 | 34 | 20 | 0 | 3.89 | 2.08 | 0 |
| | ndhU | Q84VQ4 | 24422 | 569 | 407 | 0 | 13 | 8 | 0 | 1.99 | 1.07 | 0 |
| | pnsL1 | O80634 | 26946 | 780 | 770 | 0 | 20 | 18 | 0 | 4.24 | 4.24 | 0 |
| | pnsB1 | Q9S9N6 | 50990 | 1027 | 831 | 80 | 29 | 22 | 1 | 2.08 | 1.58 | 0.06 |
| | PIFI | B3H4M0 | 23974 | 32 | 0 | 0 | 1 | 0 | 0 | 0.13 | 0 | 0 |
| **Band II** | **PSI** | | | | | | | | | | | |
| | psaA | P56766 | 83178 | 705 | 850 | 728 | 19 | 35 | 24 | 0.79 | 1.07 | 0.93 |
| | psaB | P56767 | 82423 | 642 | 1102 | 734 | 14 | 34 | 16 | 0.5 | 1.01 | 0.61 |
| | **Lhca** | | | | | | | | | | | |
| | lhca1 | A8MS75 | 23247 | 227 | 226 | 210 | 5 | 7 | 5 | 0.67 | 0.67 | 0.67 |
| | lhca2 | Q9SYW8 | 27737 | 64 | 80 | 38 | 2 | 2 | 1 | 0.11 | 0.11 | 0.11 |
| | lhca3 | Q9SY97 | 29163 | 271 | 311 | 258 | 6 | 7 | 6 | 0.67 | 0.67 | 0.67 |
| | lhca4 | P27521 | 27716 | 306 | 375 | 280 | 6 | 10 | 8 | 0.71 | 1.12 | 0.9 |
| | lhca5 | Q9C639 | 27784 | 63 | 234 | 0 | 2 | 7 | 0 | 0.24 | 1.12 | 0 |
| | lhca6 | Q8LCQ4 | 29920 | 81 | 193 | 66 | 1 | 4 | 1 | 0.1 | 0.49 | 0.1 |
| | **NDH** | | | | | | | | | | | |
| | ndhF | P56752 | 85183 | 213 | 633 | 28 | 6 | 16 | 1 | 0.84 | 1.36 | 1.36 |
| | ndhH | P56753 | 45473 | 0 | 0 | 0 | 0 | 0 | 0 | 0 | 0 | 0 |
| | ndhU | Q84VQ4 | 24422 | 0 | 0 | 0 | 0 | 0 | 0 | 0 | 0 | 0 |
| | pnsL1 | O80634 | 26946 | 158 | 702 | 0 | 4 | 17 | 0 | 0.55 | 3.68 | 0 |
| | pnsB1 | Q9S9N6 | 50990 | 281 | 615 | 97 | 7 | 18 | 3 | 0.51 | 1.03 | 0.19 |
| | PIFI | B3H4M0 | 23974 | 0 | 0 | 0 | 0 | 0 | 0 | 0 | 0 | 0 |
| **Band III** | **PSI** | | | | | | | | | | | |
| | psaA | P56766 | 83178 | 1196 | 1166 | 1215 | 66 | 101 | 68 | 1.58 | 1.49 | 1.58 |
| | psaB | P56767 | 82423 | 1276 | 1424 | 1240 | 67 | 111 | 60 | 1.16 | 1.25 | 1.08 |
| | **Lhca** | | | | | | | | | | | |
| | lhca1 | A8MS75 | 23247 | 70 | 0 | 66 | 2 | 0 | 2 | 0.29 | 0 | 0.29 |
| | lhca2 | Q9SYW8 | 27737 | 0 | 0 | 0 | 0 | 0 | 0 | 0 | 0 | 0 |
| | lhca3 | Q9SY97 | 29163 | 0 | 0 | 0 | 0 | 0 | 0 | 0 | 0 | 0 |
| | lhca4 | P27521 | 27716 | 206 | 75 | 103 | 6 | 1 | 2 | 0.71 | 0.11 | 0.24 |
| | lhca5 | Q9C639 | 27784 | 30 | 0 | 36 | 1 | 0 | 1 | 0.11 | 0 | 0.11 |
| | lhca6 | Q8LCQ4 | 29920 | 0 | 0 | 0 | 0 | 0 | 0 | 0 | 0 | 0 |
| | **NDH** | | | | | | | | | | | |
| | ndhF | P56752 | 85183 | 0 | 0 | 0 | 0 | 0 | 0 | 2.02 | 2.02 | 1.66 |
| | ndhH | P56753 | 45473 | 0 | 0 | 0 | 0 | 0 | 0 | 0 | 0 | 0 |
| | ndhU | Q84VQ4 | 24422 | 0 | 0 | 0 | 0 | 0 | 0 | 0 | 0 | 0 |
| | pnsL1 | O80634 | 26946 | 27 | 0 | 0 | 1 | 0 | 0 | 0.12 | 0 | 0 |
| | pnsB1 | Q9S9N6 | 50990 | 0 | 0 | 0 | 0 | 0 | 0 | 0 | 0 | 0 |
| | PIFI | B3H4M0 | 23974 | 0 | 0 | 0 | 0 | 0 | 0 | 0 | 0 | 0 |
